# Fatty acid desaturation by stearoyl-CoA desaturase-1 controls regulatory T cell differentiation and autoimmunity

**DOI:** 10.1101/2022.06.16.496386

**Authors:** Elien Grajchen, Melanie Loix, Paulien Baeten, Beatriz F. Côrte-Real, Ibrahim Hamad, Mansour Haidar, Jonas Dehairs, Jelle Y. Broos, James M. Ntambi, Robert Zimmermann, Rolf Breinbauer, Piet Stinissen, Niels Hellings, Gijs Kooij, Martin Giera, Johannes V. Swinnen, Bieke Broux, Markus Kleinewietfeld, Jerome J.A. Hendriks, Jeroen F.J. Bogie

## Abstract

The imbalance between pathogenic and protective T cell subsets is a cardinal feature of autoimmune disorders such as multiple sclerosis (MS). Emerging evidence indicates that endogenous and dietary-induced changes in fatty acid metabolism have a major impact on both T cell fate and autoimmunity. To date, however, the molecular mechanisms that underlie the impact of fatty acid metabolism on T cell physiology and autoimmunity remain poorly understood. Here, we report that stearoyl-CoA desaturase-1 (SCD1), an enzyme essential for the desaturation of fatty acids and highly regulated by dietary factors, acts as an endogenous brake on regulatory T cell (Treg) differentiation and augments autoimmunity in an animal model of MS. Guided by RNA sequencing and lipidomics analysis, we found that absence of *Scd1* promotes hydrolysis of triglycerides and phosphatidylcholine through adipose triglyceride lipase (ATGL). ATGL-dependent release of docosahexaenoic acid enhanced Treg differentiation by activating the nuclear receptor peroxisome proliferator-activated receptor gamma. Our findings identify fatty acid desaturation by SCD1 as an essential determinant of Treg differentiation and autoimmunity, with potentially broad implications for the development of novel therapeutic strategies and dietary interventions for autoimmune disorders.

## Introduction

Autoimmunity reflects an imbalance between effector and regulatory mechanisms, resulting in loss of immunological self-tolerance. Multiple sclerosis (MS) is a devastating neurological disease and one of the most prevalent autoimmune disorders in the Western world. The autoimmune response in MS is characterized by an increase in autoreactive pro-inflammatory immune cell subsets, *i*.*e*. T helper (Th) 17 and Th1 cells, and a decrease in number and function of regulatory T cells (Tregs) (1-3). This imbalance between pathogenic and protective T cell subsets is considered to drive demyelination and neurodegeneration in the central nervous system (CNS). Hence, restoring the immune balance in favor of protective T cell subsets represents a promising strategy to halt MS disease progression.

Emerging evidence indicates that fatty acids control adaptive immunity and MS disease pathology (4). Early epidemiological studies found a strong association between excessive fat intake, obesity, and the etiology of MS (5, 6). Notwithstanding, the direct effects of dietary fatty acids on immune cell function and CNS pathology are only recently being uncovered. Carbon chain length and degree of desaturation appear to be key determinants that dictate the immunopathological outcome of dietary fatty acids in MS, with saturated short-chain, monounsaturated long-chain, and polyunsaturated very long-chain fatty acids inducing protective immunity and reducing CNS pathology (7-10). Several recent studies also defined the importance of endogenous fatty acid metabolism pathways in immune cell physiology. For example, the formation of specialized pro-resolving lipid mediators was found to be essential for inducing a protective adaptive immune response (11, 12). Furthermore, inhibition of *de novo* fatty acid synthesis and stimulation of beta-oxidation attenuate neuroinflammation and favor the differentiation of Tregs at the expense of Th17 cells (13-18). Here, changes in the metabolism and intracellular levels of oleic (C18:1) and palmitic acid (C16:0) triggered a neuroprotective adaptive immune response (13, 18).

In this study, we defined the importance of stearoyl-CoA desaturase-1 (SCD1), an enzyme that catalyzes the rate-limiting step in the conversion of saturated fatty acids (SFAs, C16:0 and C18:0) into mono-unsaturated fatty acids (MUFAs, C16:1 and C18:1), in T cell physiology and neuroinflammation. SCD1 belongs to the family of Δ9 fatty acid desaturases whose expression and activity is highly responsive to dietary factors, being negatively and positively regulated by polyunsaturated fatty acids (PUFAs) and Western-type diets rich in SFAs, respectively (4, 19). We show that pharmacological inhibition of SCD1 and *Scd1* deficiency doubles peripheral Treg numbers and attenuates disease severity in experimental autoimmune encephalomyelitis (EAE), the most commonly used animal model to study MS. Guided by RNA sequencing and lipidomics analyses, we reveal that *Scd1* deficiency promotes Treg differentiation through adipose triglyceride lipase (ATGL)-dependent hydrolysis of triglycerides and phosphatidylcholine, thereby releasing non-esterified bioactive docosahexaenoic acid (DHA, C22:6), a natural ligand of the nuclear receptor peroxisome proliferator-activated receptor gamma (PPARγ). Our findings highlight the importance of SCD1 in controlling Treg differentiation and reveal its potential as a therapeutic target for MS and other autoimmune diseases.

## Results

### SCD1 deficiency and pharmacological inhibition reduces EAE severity by promoting Treg differentiation

Fatty acid desaturation and dietary factors are closely associated with MS disease progression (5-7, 9). Given its essential role in controlling fatty acid desaturation and its expression being highly susceptive to dietary factors, we defined the impact of SCD1 on the disease severity and immunopathology of EAE. By using a mouse strain deficient in *Scd1* (*Scd1*^-/-^), we found that absence of *Scd1* attenuated EAE disease severity and incidence, and significantly delayed disease onset (Figure 1A and Supplemental Figure 1, A and B). Pathologically, *Scd1*^-/-^ mice showed a decreased inflammatory expression profile as compared to wt animals, both at 18 days post-immunization and their respective disease peaks (Figure 1B). Furthermore, a reduced infiltration of CD3^+^ T cells and F4/80^+^ phagocytes was observed in the spinal cord of *Scd1*^*-/-*^ EAE mice at 18 dpi (Figure 1, C and D). Consistent with *Scd1* deficiency, animals treated with a pharmacological inhibitor of SCD1 (CAY10566) showed reduced EAE disease severity and incidence, delayed disease onset, and pathological parameters largely matched those observed in *Scd1*^-/-^ mice (Figure 1, E-H, and Supplemental Figure 1, C and D).

**Figure 1.**
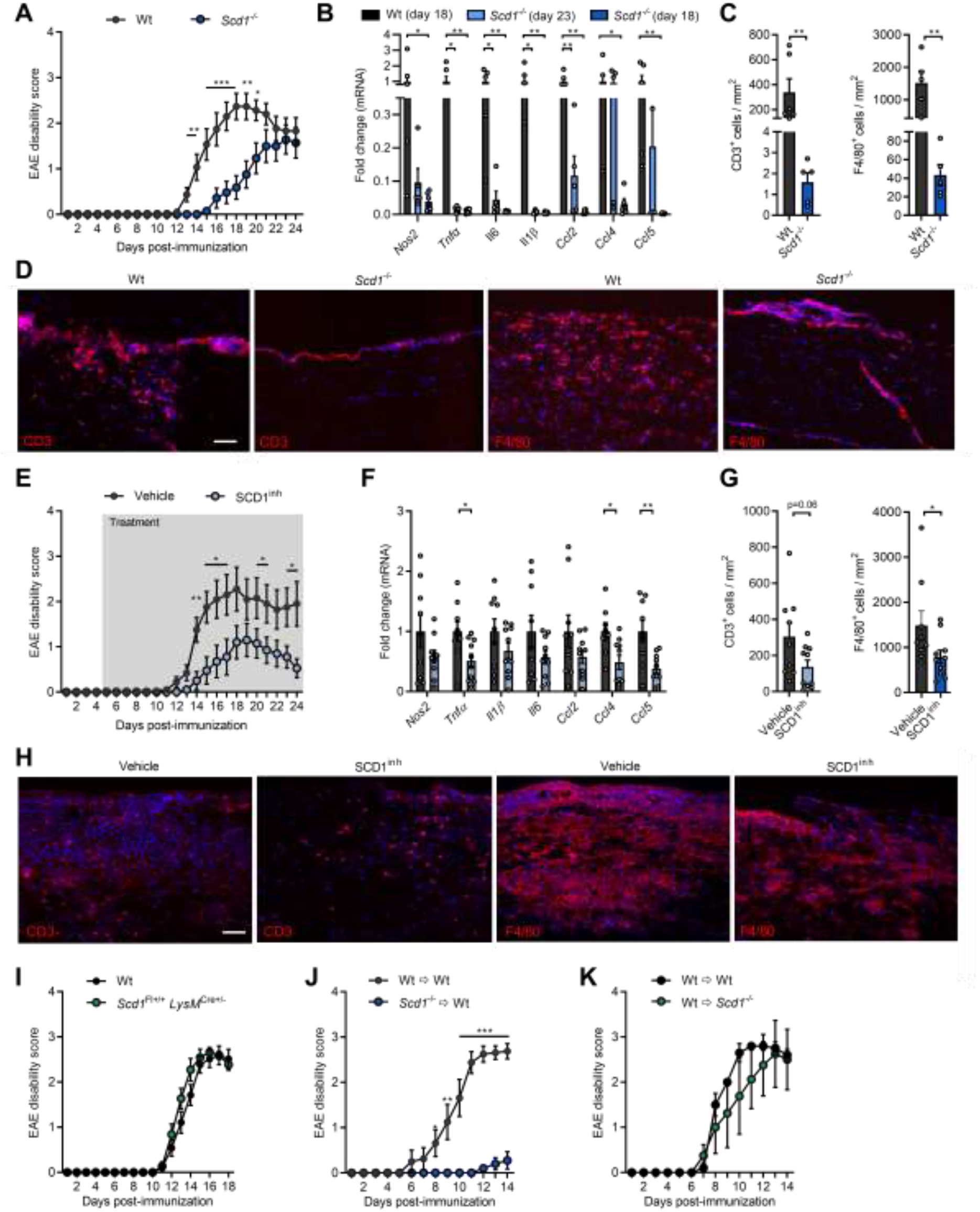
SCD1 inhibition and deficiency reduces EAE severity in a T cell-dependent manner. **(A)** EAE disease score of *Scd1*^-/-^ mice (n=14) and wild-type littermates (wt, n=15). **(B)** mRNA expression of *Nos2, Tnfα, Il1β, Il6, Ccl2, Ccl4*, and *Ccl5* in spinal cord tissue obtained from wt (n=6) and *Scd1*^-/-^ (n=5) EAE animals (18 and 23 days post-immunization, dpi). **(C**,**D)** Quantification (C) and representative images (D) of CD3 and F4/80 staining of spinal cord tissue obtained from wt (n=6) and *Scd1*^-/-^ (n=5) EAE animals at the peak of the disease (18 dpi). Scale bar: 100 µm. **(E)** EAE disease score of wt mice treated with vehicle (n=10) or SCD1 inhibitor (SCD1^inh^, CAY10566, 2.5 mg/kg, n=10) by oral gavage twice a day with a 12 h interval. **(F)** mRNA expression of *Nos2, Tnfα, Il1β, Il6, Ccl2, Ccl4*, and *Ccl5* in spinal cord tissue obtained from vehicle-(n=9) and SCD1 inhibitor-treated (n=10) EAE animals (24 dpi). **(G**,**H)** Quantification (G) and representative images (H) of CD3 and F4/80 staining of spinal cord tissue obtained from vehicle-(n=9) and SCD1 inhibitor-treated (n=10) EAE animals (24 dpi). **(I)** EAE disease score of wt (*Scd1*^Fl+/+^ and *LysM*^Cre+/-^, n=19) and *Scd1*^Fl+/+^ *LysM*^Cre+/-^ (n=11) mice. **(J)** EAE disease score of wt recipient mice that received lymph node-derived T lymphocytes from immunized wt (Wt ➩ Wt, n=8) or *Scd1*^-/-^ mice (*Scd1*^-/-^ ➩ Wt, n=10). **(K)** EAE disease score of wt (n=5) and *Scd1*^-/-^ (n=4) recipient mice that received lymph node-derived T lymphocytes from immunized wt animals. All replicates were biologically independent. (A,E,J) Statistics denote differences between wt and *Scd1*^-/-^ at the time points indicated. All data are represented as mean ± SEM. *, P < 0.05; **, P < 0.01; ***, P < 0.001; calculated with Tukey’s post hoc analysis (B) or a two-tailed unpaired student T-test (A,C,E-K).

Previously, we demonstrated that *Scd1* deficiency induces a less-inflammatory, repair-promoting phenotype in macrophages and microglia in the CNS (20). To assess whether these cells are responsible for reduced EAE disease severity in *Scd1*^-/-^ mice, EAE was induced in mice with macrophage/microglia-specific *Scd1* deficiency (*Scd1*^Fl+/+^ *LysM*^Cre+/-^). Interestingly, we found no difference in EAE disease severity, incidence, and onset between wt and *Scd1*^Fl+/+^ *LysM*^Cre+/-^ mice (Figure 1I and Supplemental Figure 1, E and F). These findings indicate that attenuation of EAE symptoms by *Scd1* deficiency is not mediated by macrophages and microglia. To assess whether T cells are responsible for reduced EAE severity in *Scd1*^-/-^ mice, we applied an adoptive transfer EAE model. For this purpose, lymph node-derived T lymphocytes were transferred from immunized wt or *Scd1*^-/-^ mice to wt recipient mice. Strikingly, animals that received *Scd1*^-/-^ T cells showed almost no EAE symptoms compared to those transplanted with wt T cells (Figure 1J and Supplemental Figure 1, G and H). Accordingly, we observed no difference in EAE disease severity, incidence, and onset after transfer of wt T cells from immunized mice to wt or *Scd1*^-/-^ mice (Figure 1K and Supplemental Figure 1, I and J). Altogether, these findings indicate that absence of SCD1 ameliorates EAE disease severity in a T cell-dependent manner.

To define the culprit T cell subset, we characterized the peripheral T cell compartment of wt and *Scd1*^-/-^ EAE animals prior to disease onset. No differences were detected in the absolute number of lymph node immune cells between wt and *Scd1*^-/-^ animals (data not shown), nor in the relative abundance of CD8^+^ and CD4^+^ T cells (Figure 2, A and B). Similar, the frequency of the T helper subsets, *i*.*e*. CD4^+^IFNγ^+^ Th1 cells, CD4^+^IL17^+^ Th17 cells, and CD4^+^IL4^+^ Th2 cells, remained unaffected in *Scd1*^-/-^ mice (Figure 2, A and B). Interestingly, we detected a remarkable two-fold higher frequency of CD4^+^FOXP3^+^ Tregs in the lymph nodes of *Scd1*^-/-^ mice (Figure 2, A and B). Similar findings were observed in the spleen of wt and *Scd1*^-/-^ EAE animals (Figure 2C and Supplemental Figure 2A). To confirm the essential role of SCD1 in Treg physiology, we next assessed the impact of SCD1 on Treg differentiation in vitro. We observed that both genetic deficiency and pharmacological inhibition of SCD1 promotes the differentiation of Tregs from murine naïve T cells (Figure 2D and Supplemental Figure 2B). A similar increase in Treg differentiation was observed using human PBMC-derived naïve T cells after exposure to the SCD1 inhibitor (Figure 2E and Supplemental Figure 2C). Of note, absence or inhibition of SCD1 did not impact the suppressive capacity of mouse and human Tregs (Figure 2, F and G, and Supplemental Figure 3, A and B), nor did it increase Treg proliferation (Supplemental Figure 3C). Collectively, these findings show that SCD1 acts as a brake on the differentiation of Tregs.

**Figure 2.**
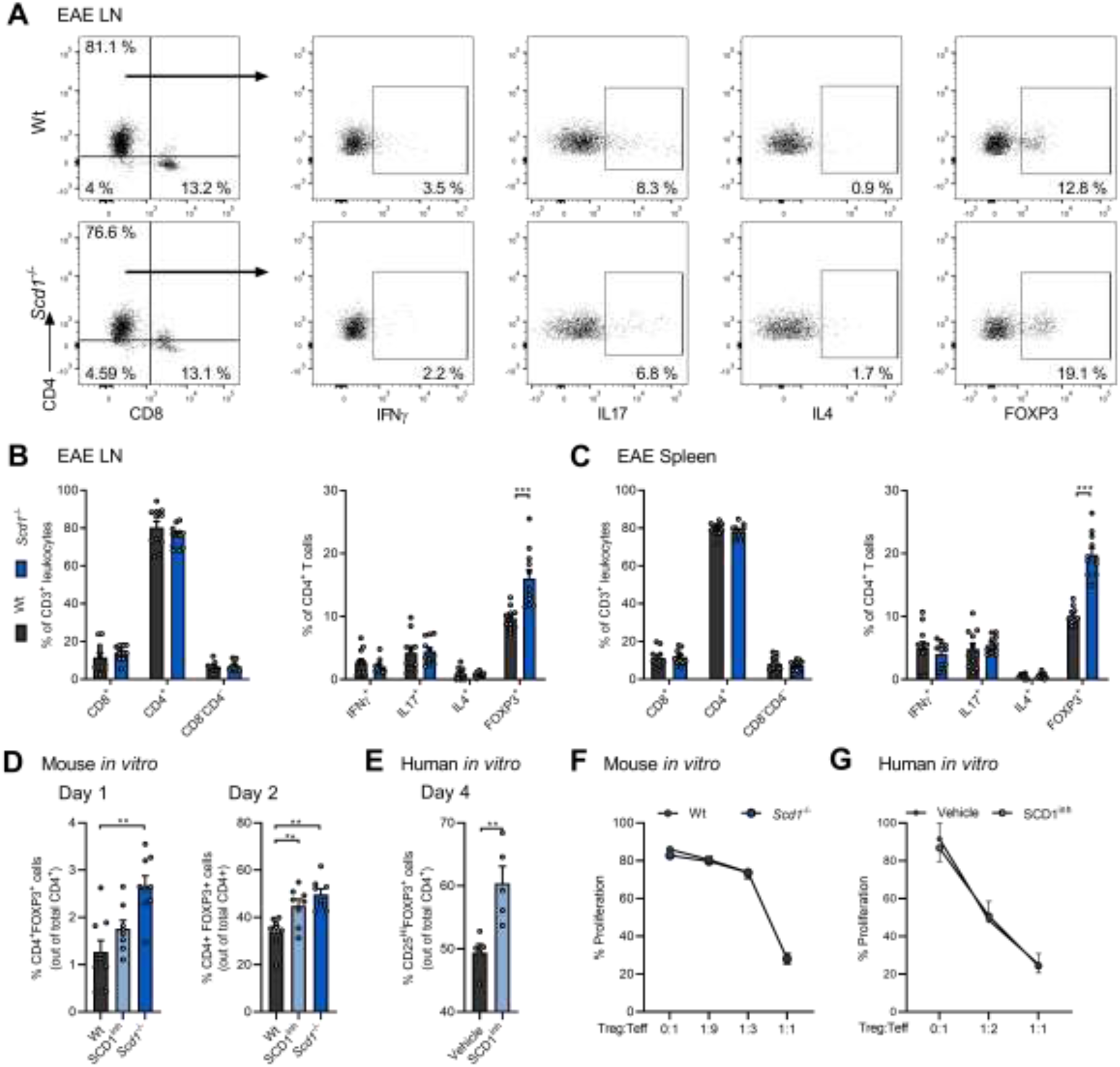
SCD1 deficiency and inhibition promotes the differentiation of Tregs. **(A-C)**Frequency of CD4^+^, CD8^+^, CD4^-^CD8^-^, CD4^+^IFNγ^+^, CD4^+^IL17^+^, CD4^+^IL4^+^, and CD4^+^FOXP3^+^ cells in the lymph nodes (LN, A and B) and spleen (C) of wild-type (wt, n=11 animals) and *Scd1*^-/-^ (n=11 animals) EAE animals 10 days post-immunization (dpi). Representative flow cytometric plots (A) and flow cytometric analysis (B,C) are shown. **(D**,**E)** Wt and *Scd1*^-/-^ mouse naïve T cells (D) and human naïve T cells (E) were differentiated under Treg polarizing conditions and treated with vehicle or SCD1 inhibitor (SCD1^inh^, CAY10566, 1 µM). For mouse T cell cultures, the frequency of CD4^+^FOXP3^+^ cells was quantified one and two days after Treg induction (n=8 samples). For human T cell cultures, frequency of CD25^Hi^FOXP3^+^ cells was quantified four days after induction (n=5 healthy controls). Results from (A-E) are pooled from at least three independent experiments. **(F**,**G)** Suppressive capacity of wt and *Scd1*^*-/-*^ mouse Tregs (F), or human Tregs differentiated in the presence of an SCD1 inhibitor (SCD1inh, CAY10566, 1 µM) or vehicle (G). Increasing amounts of Tregs were cultured with CFSE- or CellTrace Violet (CTV)-labeled CD4^+^CD25^-^ effector T cells (F, n=3 wt samples; G, n=2 healthy controls). Percentage proliferation was assessed after three (F) or five (G) days. All data are represented as mean ± SEM. *, P < 0.05; **, P < 0.01; ***, P < 0.001, calculated with two-tailed unpaired student T-test (B,C), Tukey’s post hoc analysis (D), or Mann-Whitney analysis (E-G).

### *Scd1*-deficient naïve T cells display a transcriptional profile characteristic of Tregs

To identify transcriptional alterations that underlie the impact of *Scd1* deficiency on T cell physiology, bulk RNA sequencing was performed. Differential gene expression analysis revealed that 235 genes distinguished wt and *Scd1*^-/-^ naïve CD4^+^ T cells, with 135 more highly expressed in *Scd1*^-/-^ naïve T cells and 100 more highly expressed in wt cells (Figure 3A and Supplemental Table 2). By using Ingenuity Pathway Analysis, we found that differentially expressed genes were highly associated with cancer and tumorigenic processes (Supplemental Figure 4A), corresponding to the key role of SCD1 in cancer cell migration, metastasis, and tumor growth (21). Canonical pathway analysis identified 34 enriched biological pathways (Supplemental Figure 4B), including ‘DHA signaling’, ‘unfolded protein response’, and ‘amyotrophic lateral sclerosis signaling’ (Figure 3, B and C). Molecular and cellular functions that matched the transcriptional profile of naïve *Scd1*^-/-^ T cells were mapped to phospholipid, cholesterol, and acylglycerol metabolism (Figure 3, D and E). These findings confirm the essential role of SCD1 in controlling cellular lipid metabolism.

**Figure 3.**
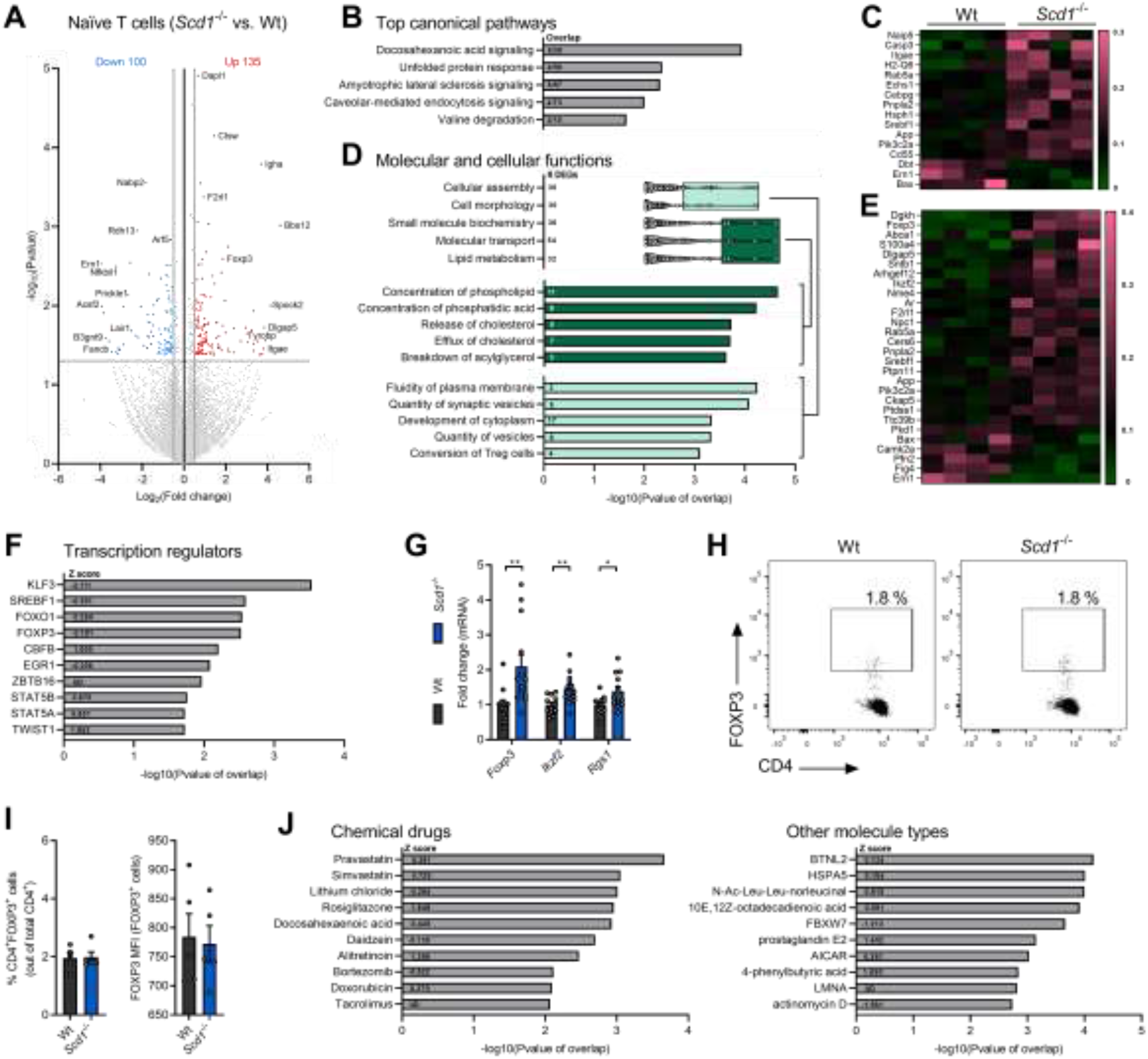
Naïve *Scd1*^-/-^ T cells display a transcriptional profile characteristic of Tregs. (A-F, J) Bulk RNA sequencing was performed using wild-type (wt) and *Scd1*^-/-^ naïve CD4^+^ T cells (CD4^+^CD25^-^CD44medCD62L^+^). **(A)** Differential gene expression analysis of the RNA sequencing data (log _2_ fold change < -0.5 and >0.5; P value <0.05; complete list in Supplementary Table 2). **(B)** Top five most enriched canonical pathways in *Scd1*^-/-^ naïve T cells (complete list in Supplemental Figure 3b). **(C)** Heat map displaying the expression of genes present in the top five canonical pathways. The data was scaled by the sum of each row. **(D)** Molecular and cellular function categories associated with differentially expressed genes in *Scd1*^-/-^ naïve T cells. **(E)** Heat map displaying the expression of genes involved in the molecular and cellular function categories. The data was scaled by the sum of each row. **(F)** Top 10 transcriptional regulators most associated with the transcriptome of *Scd1*^-/-^ naïve T cells. **(G)** mRNA expression of *Foxp3, Ikzf2*, and *Rgs1* in wt and *Scd1*^-/-^ naïve T cells (n=11 samples). **(H, I)** Frequency of CD4^+^FOXP3^+^ cells and respective mean fluorescence intensity of FOXP3 in wt and *Scd1*^-/-^ naïve T cell population was quantified (n=5 samples). Representative flow cytometric plots (H) and flow cytometric analysis (I) are shown. **(J)** Top 10 chemical drugs, transcriptional regulators, and other molecule types most associated with the transcriptome of *Scd1*^-/-^ naïve T cells. Results are pooled from four (A-F, I and J), seven (G), or five (H) independent experiments. Data from (G,I) are represented as mean ± SEM. *, P < 0.05; **, P < 0.01, calculated with two-tailed unpaired student T-test (G,I).

Interestingly, ‘conversion of Treg cells’ was recognized as a key cellular process in naïve *Scd1*^-/-^ T cells (Figure 3, D and E). Accordingly, based on increased expression of *Foxp3, Ikzf2*, and *Rgs1*, upstream analysis demonstrated Treg-associated transcription factors FOXO1, FOXP3, CBFB, and STAT5 as significant upstream regulators of the transcriptional signature of naïve *Scd1*^-/-^ T cells (22-25) (Figure 3, E and F). Elevated expression of Treg-associated genes *Foxp3, Ikzf2*, and *Rgs1* was validated using qPCR (Figure 3G). In contrast to naïve T cells, *Scd1*-deficient Tregs did not show an altered expression of *Foxp3, Ikzf2*, and *Rgs1*, nor did they demonstrate changes in the expression of metabolic genes that were differentially regulated in naïve wt and *Scd1*^*-/-*^ T cells (Supplemental Figure 5). Of interest, flow cytometric analysis demonstrated no difference in the frequency of FOXP3^+^ cells, nor in FOXP3 protein abundance between wt and *Scd1*^-/-^ naïve T cells (Figure 3, H and I). These findings suggest that *Scd1* deficiency induces a transcriptional profile characteristic of Tregs in naïve T cells, without affecting the transcriptome of differentiated committed Tregs.

Chemical drugs and other molecule types associated with the transcriptome of naïve *Scd1*^-/-^ T cells included cholesterol-reducing agents (pravastatin and simvastatin) and activators of the nuclear receptor PPARγ (rosiglitazone, docosahexaenoic acid, 10E, 12Z-octadecadienoic acid, prostaglandin E2, and 4-phenylbutyric acid) (Figure 3J). The positive association between these lipid-modifying chemical drugs and fatty-acid containing lipid species confirms the pathway and function enrichment analysis, and further underlines the role of SCD1 in cellular lipid metabolism. In conclusion, pathway and gene set enrichment analysis indicate that *Scd1* deficiency in naïve T cells affects DHA and PPARγ signaling, alters phospholipid and triglyceride metabolism, and induces a transcriptional profile characteristic of Tregs.

### *Scd1* deficiency results in PUFA-depleted lysolecithin and triglycerides

Given the overrepresentation of differentially expressed genes in pathways related to lipid metabolism, liquid chromatography electrospray ionization tandem mass spectrometry (LC-ESI-MS/MS) analysis was performed to define the fatty acid lipidome of wt and *Scd1*^-/-^ naïve T cells. Consistent with reduced SCD1 activity, naïve *Scd1*^-/-^ T cells demonstrated decreased 16:1/16:0 and C18:1/C18:0 desaturation indices, particularly within phospholipid classes (Supplemental Figure 6, A and B). Of interest, naïve *Scd1*^-/-^ T cells showed decreased levels of all major lipid classes except for lysophosphatidylcholine (LPC), a hydrolysed derivative of phosphatidylcholine (PC; Figure 4, A and B, detailed information in Supplemental Figure 6, C-R), which argues for elevated hydrolysis of PC in naïve *Scd1*^-/-^ T cells. The overall decrease in cellular lipids in naïve *Scd1*^-/-^ T cells did not depend on reduced cell size or granularity (Supplemental Figure 6S).

**Figure 4.**
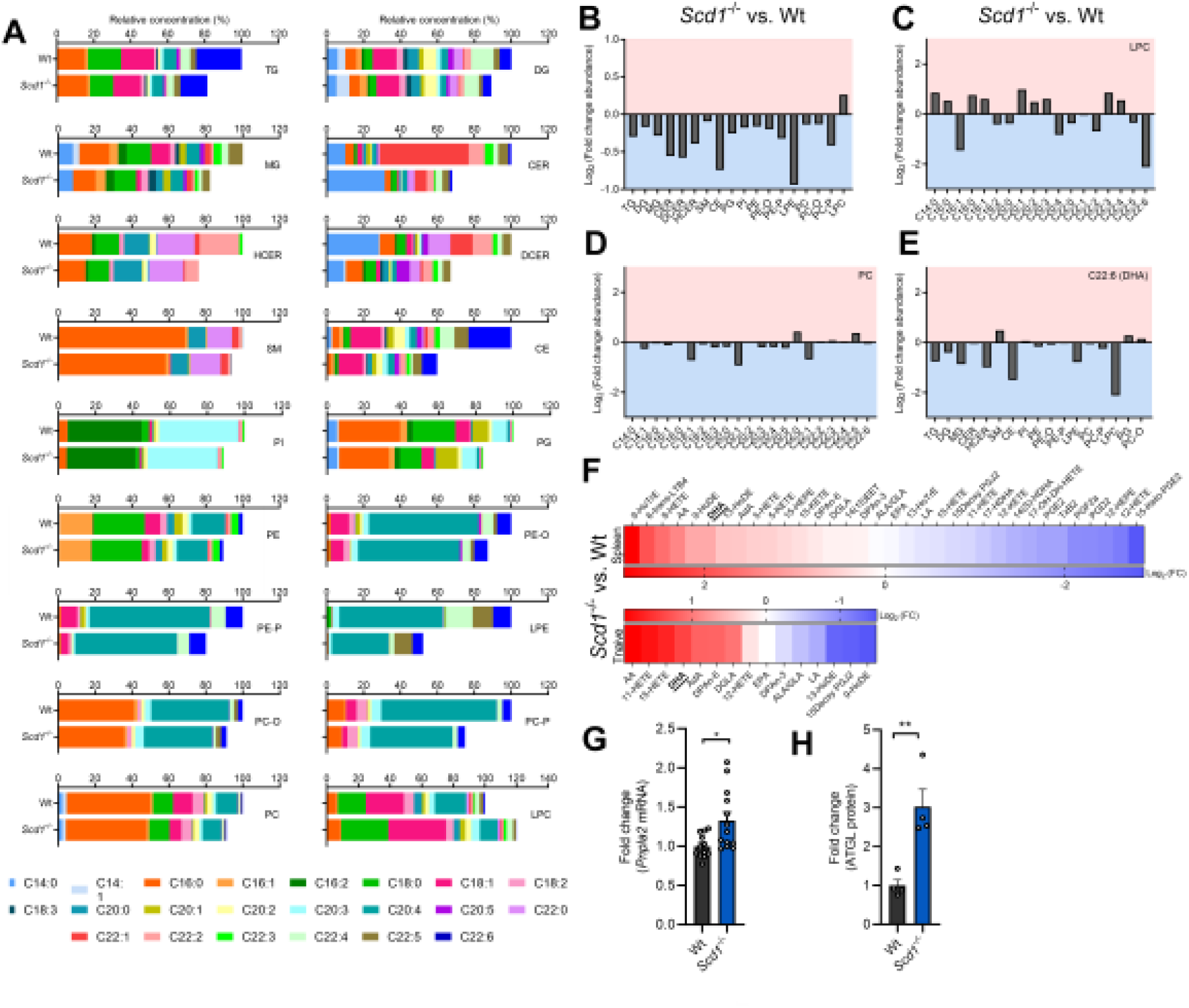
*Scd1* deficiency results in DHA-depleted lysolecithin and triglycerides. **(A-E)** Liquid chromatography electrospray ionization tandem mass spectrometry (LC-ESI-MS/MS) analysis was performed to define the lipidome of wild-type (wt) and *Scd1*^-/-^ naïve T cells (n=2 samples). **(A)** Fatty acid composition of all lipid classes. **(B-E)** Log _2_ fold change abundance of all lipid classes (B), fatty acyl moieties within lysophosphatidylcholine (LPC, C) and phosphatidylcholine (PC, D), and docosahexaenoic acid (DHA) within all lipid classes (E) is shown (*Scd1*^-/-^ vs. wt). Only detectable fatty acyl moieties and non-esterified fatty acids and downstream metabolites are reported. **(F)** LC-MS/MS analysis was performed to determine the abundance of non-esterified fatty acids and downstream metabolites in wt and *Scd1*^-/-^ naïve T cells and spleen tissue (n=3 samples). Log _2_ fold change abundance of all fatty acids and downstream metabolites is shown (*Scd1*^-/-^ vs. wt). **(G)** mRNA expression of *Pnpla2* in wt and *Scd1*^-/-^ naïve T cells (n=11 samples). **(H)** ATGL protein level in wt and *Scd1*^-/-^ naïve T cells (n=4 samples). Results are pooled from two (A-E, H), three (F), or seven (G) independent experiments. Data are represented as mean (A-F) or as mean ± SEM (G). *, P < 0.05; **, P<0.01, calculated with two-tailed unpaired student T-test (G,H).

In-depth analysis further demonstrated that LPC in naïve *Scd1*^-/-^ T cells was mainly depleted of the ω3 PUFA DHA (C22:6), aside from the expected decrease in palmitoleic acid (C16:1) (Figure 4C). This decrease of DHA in LPC was not apparent in PC (Figure 4D). Interestingly, mono-, di-, and triacylglycerides (MGs, DGs, and TGs) showed a marked decrease in DHA as well (Figure 4E), mirroring the enrichment of genes involved in the catabolism of acylglycerols in *Scd1*^-/-^ naïve T cells (Figure 3D). These findings point towards enhanced activity of a lipase that promotes the hydrolysis of PC and acylglycerols, and favors the release of DHA in *Scd1*^-/-^ naïve T cells. In support of the latter, LC-MS/MS analysis revealed an increased abundance of non-esterified PUFAs, including DHA, in spleen tissue and naïve T cells from *Scd1*^-/-^ animals (Figure 4F). To identify the lipase involved, we re-evaluated differentially expressed genes obtained from our transcriptomic analysis. We found that gene expression and protein abundance of *Pnpla2*, which encodes for the enzyme adipose triglyceride lipase (ATGL), was elevated in naïve *Scd1*^-/-^ T cells (Figure 3, C and E, and Figure 4, G and H). ATGL is a calcium-independent cellular phospholipase that hydrolyses TGs and phospholipids rich in DHA and arachidonic acid (AA) (26). With respect to the latter, *Scd1*^*-/-*^ T cells also showed a reduced abundance of AA in LPC and elevated levels of non-esterified AA (Figure 4, C and F). Collectively, these data suggest that *Scd1* deficiency enhances ATGL-mediated hydrolysis of PC and TGs, thereby releasing DHA and AA. Given that the release of phospholipid- and acylglycerol-associated DHA is essential for many of its biological activities as well as its capacity to activate nuclear receptors (27), the lipidomics analyses further provide a molecular rationale for the enrichment of genes in the DHA and PPARγ signaling pathways in *Scd1*^-/-^ T cells (Figure 3, B-E and J).

### ATGL-driven hydrolysis of PC and TGs stimulates Treg differentiation in *Scd1*^-/-^ T cells by releasing DHA

Based on the RNA sequencing and lipidomics analysis, we next assessed the importance of ATGL and DHA in driving enhanced Treg differentiation of naive *Scd1*^-/-^ T cells. By using a pharmacological inhibitor of ATGL (Atglistatin), we found that ATGL inhibition counteracted the enhanced induction of Tregs from *Scd1*^-/-^ naïve T cells (Figure 5, A and B), and in parallel, markedly reduced intracellular abundance of non-esterified DHA (Supplemental Figure 7A). ATGL inhibition did not decrease the capacity of wt naïve T cells to differentiate into Tregs (Figure 5, A and B). Similar findings were obtained following shRNA-mediated gene silencing of *Atgl* in mouse T cells and upon exposure of human PBMC-derived naïve T cells to the ATGL inhibitor (Figure 5, C-E, and Supplemental Figure 7B). Importantly, ATGL inhibition also decreased Treg frequency in the spleen of EAE animals exposed to the SCD1 inhibitor (Figure 5F and Supplemental Figure 7C), highlighting the essential role of the SCD1-ATGL axis in autoimmunity under physiological conditions. In support of a key role of DHA in driving Treg differentiation, exposure to DHA significantly promoted Treg differentiation (Figure 5, G and H). Together, these data indicate that enhanced Treg differentiation upon SCD1 inhibition and in *Scd1*^-/-^ T cells partially relies on increased ATGL activity. They further suggest that non-esterified bioactive DHA, released through ATGL-mediated hydrolysis of PC and acylglycerols, acts as an important lipid metabolite in this pathway.

**Figure 5.**
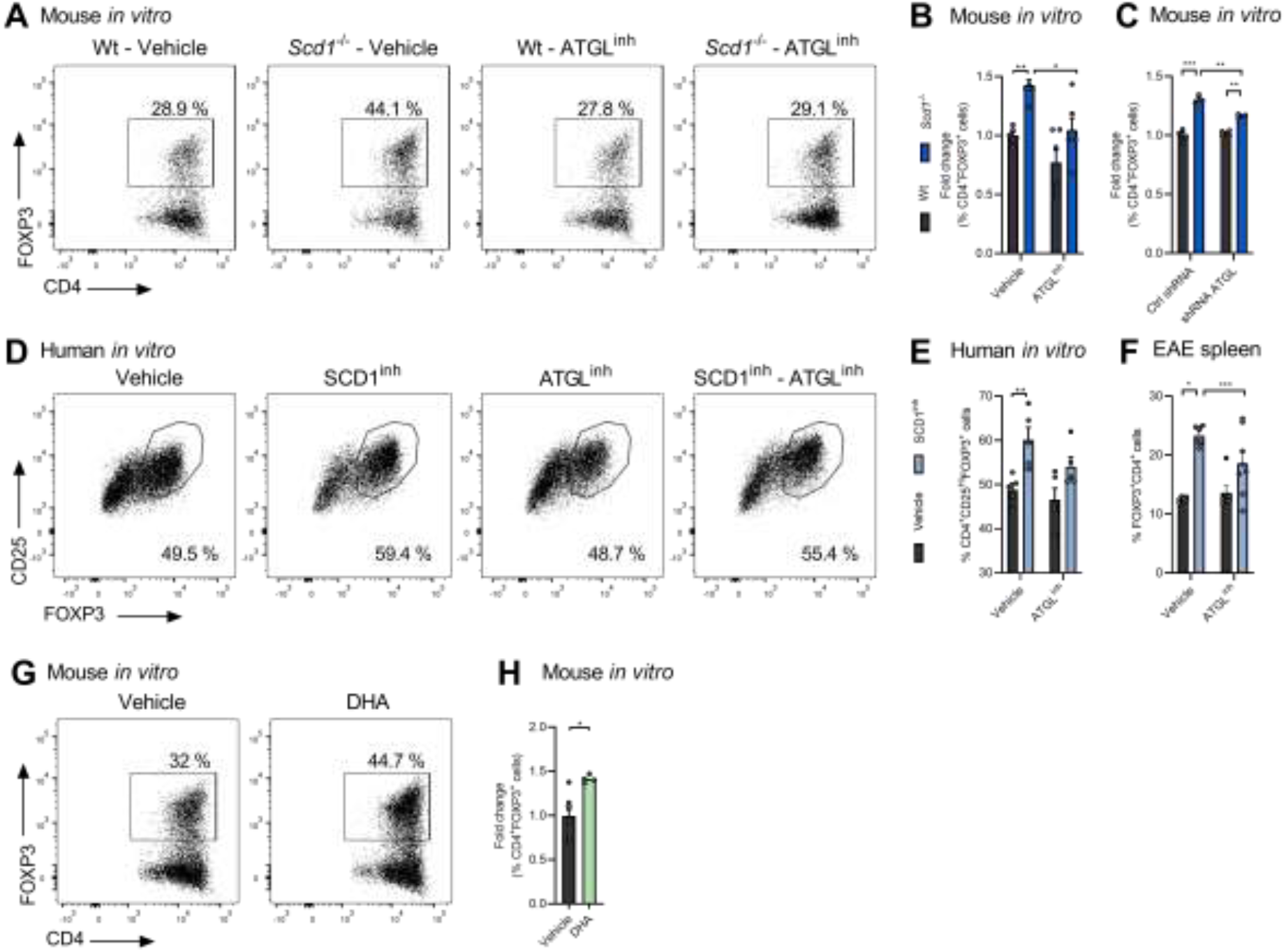
ATGL-driven release of non-esterified DHA enhances Treg differentiation in *Scd1*^-/-^ T cells. **(A-C)** Wt and *Scd1*^-/-^ mouse naïve T cells were differentiated under Treg polarizing conditions and treated with vehicle or ATGL inhibitor (ATGL^inh^, Atglistatin, 20 µM; n=9 samples), or were transduced with ATGL specific (shRNA ATGL) or control shRNA (Ctrl shRNA) (n=4 samples). The frequency of CD4^+^FOXP3^+^ cells was quantified two days after Treg induction. **(D-E)** Human naïve T cells were treated with SCD1 inhibitor (SCD1^inh^, CAY10566, 1 µM), ATGL^inh^, or vehicle (n=5 healthy controls). Frequency of CD25^Hi^FOXP3^+^ cells was quantified four days after Treg induction. Representative flow cytometric plots (A, D) and flow cytometric analyses (B, C, E) are shown. **(F)** Flow cytometric analysis of CD4^+^FOXP3^+^ cells in the spleen of EAE mice treated with an SCD1 inhibitor (CAY10566, 2.5 mg/kg) and ATGL inhibitor (Atglistatin, 200 µmol/kg) or vehicle (methylcellulose) by oral gavage twice a day with a 12 h interval, starting 5 dpi. Cells were collected 10 days post-immunization. **(G, H)** Wt and *Scd1*^-/-^ mouse naïve T cells were differentiated under Treg polarizing conditions and treated with vehicle or BSA-conjugated DHA (1 µM; n=4 samples). Results are pooled from at least two independent experiments and presented as mean ± SEM. *, P < 0.05; **, P < 0.01; ***,P <0.001, calculated with two-tailed unpaired student T-test (B,C,E,F,H).

### Non-esterified DHA promotes Treg differentiation through PPARγ

Upstream analysis revealed that differentially expressed genes in *Scd1*^-/-^ naïve T cells were associated with the activation of PPARγ (Figure 3J). Consistent with these findings, naïve *Scd1*^*-/-*^ T cells demonstrated an increased expression of PPARγ responsive genes, including *Srebpc1, Abca1, Cd36, Cpt1a*, and *Lpl* (Supplemental Figure 8A). Furthermore, by using ligand-binding luciferase reporter assays, we show that SCD1 inhibition promoted the ligation of PPARγ but not PPARα or PPARβ (Figure 6A). Interestingly, inhibition of ATGL countered the ligation of PPARγ induced by SCD1 inhibition (Figure 6A). In line with a key role of non-esterified DHA in the activation of PPARs, exposure to DHA induced the activation of PPARγ and PPARβ, but not PPARα (Figure 6B). Collectively, these findings strongly suggest that *Scd1* deficiency results in ATGL-mediated release of DHA, which in turn activates PPARγ.

Given that PPARγ activation promotes Treg differentiation and that DHA is an endogenous PPARγ ligand (27-29), we next assessed the role of PPARs in driving enhanced Treg differentiation in *Scd1*^-/-^ naïve T cells. To this end, T cell cultures were exposed to antagonists for the different PPAR isoforms. While both PPARβ and PPARγ were found to be important for Treg differentiation, only PPARγ antagonism prevented enhanced mouse Treg differentiation induced by the absence of *Scd1* (Figure 6, C and D). Consistent with these findings, *Pparγ* gene silencing counteracted enhanced Treg differentiation in *Scd1*^-/-^ T cell cultures (Figure 6E and Supplemental Figure 8B). Similar to murine T cells, treatment with the PPARγ antagonist reduced Treg differentiation independently of SCD1 and abolished enhanced Treg differentiation caused by SCD1 inhibition in human T cell cultures (Figure 6F and Supplemental Figure 8C).

**Figure 6.**
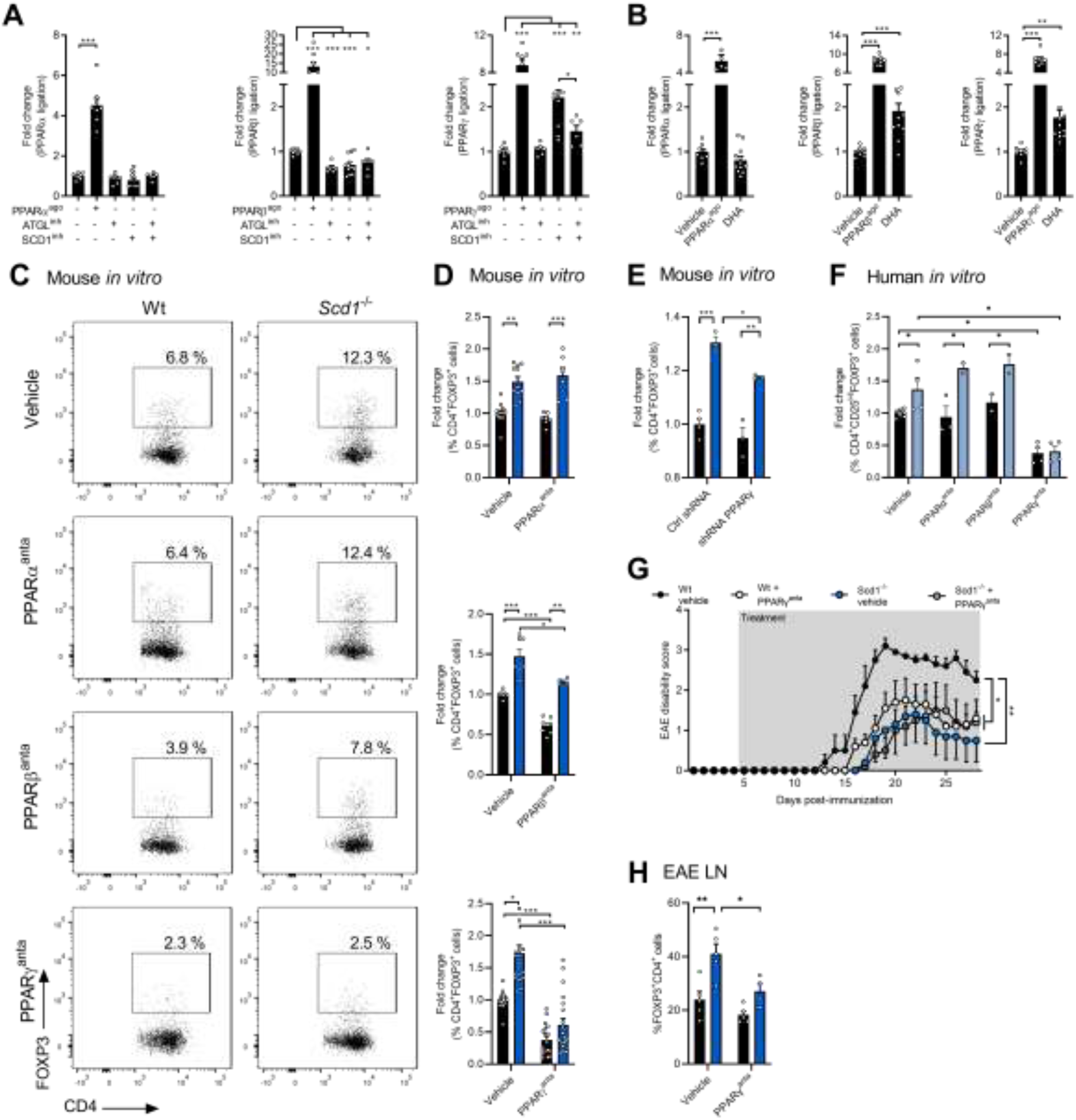
DHA-PPARγ signaling promotes Treg differentiation in the absence of SCD1. **(A**,**B)** Ligand-binding luciferase reporter assays were used to assess ligation of the different PPAR isoforms (n=6-12 samples). Jurkat T cells were treated with vehicle, ATGL inhibitor (ATGLinh, atglistatin, 20 μM), SCD1 inhibitor (SCD1inh, CAY10566, 10 μM) (A), DHA (1 μM) (B), and/or PPAR agonists (PPARα^ago^: WY-14643, 10 μM; PPARβ^ago^: GW501516, 10 μM; PPARγ^ago^: rosiglitazone, 10 μM) for 24 h, after which PPAR ligation was assessed. **(C**,**D)** Wild-type (wt) and *Scd1*^-/-^ naïve T cells were differentiated under Treg polarizing conditions and treated with vehicle, PPARα antagonist (PPARα^anta^: GW6471, 25 μM; n=7-13 samples), PPARβ antagonist (PPARβ^anta^: PTS58, 25 μM; n=5-7 samples), and PPARγ antagonist (PPARγ^anta^: GW9662, 25 μM; n=18-20 samples). Frequency of CD4^+^FOXP3^+^cells was quantified one day after Treg induction. Representative flow cytometric plots (C) and flow cytometric analysis (D) are shown. **(E)** Wt and *Scd1*^-/-^ naïve T cells were transduced with PPARγ specific shRNA (shRNA PPARγ) or control shRNA (Ctrl shRNA), followed by Treg differentiation and analysis. **(F)** Human naïve T cells were differentiated under Treg polarizing conditions and treated with vehicle (n=5 healthy controls), PPARα antagonist (PPARα^anta^: GW6471, 25 μM; n=3 healthy controls), PPARβ antagonist (PPARβ^anta^: PTS58, 25 μM; n=3 healthy controls) or PPARγ antagonist (PPARγ^anta^: 25 μM; n=4 healthy controls). Frequency of CD25^+^FOXP3^+^ cells of total CD4^+^ cells was quantified four days after Treg induction by flow cytometry. **(G**,**H)** Wt and *Scd1*^*-/-*^ EAE mice were treated daily with vehicle or a PPARγ-antagonist (GW9662, 2mg/kg, n=5/group). Lymph nodes were collected 28 days post-immunization. EAE disease score (G) and flow cytometric analysis of frequency of CD4^+^FOXP3^+^ cells in the lymph nodes (H). Data are represented as mean ± SEM. **, P < 0.01, calculated with Tukey’s post hoc analysis.

Finally, to confirm the interplay between SCD1 and PPARγ in autoimmunity under physiological conditions, *Scd1*^-/-^ EAE animals were exposed to the PPARγ antagonist GW9662. Here, we observed that *Scd1* deficiency no longer reduced EAE disease severity in the presence of GW9662 as compared to control animals treated with GW9662 (Figure 6G). Surprisingly, PPARγ antagonism reduced EAE disease severity as compared to control mice (Figure 6G). The latter finding contradicts with the protective impact of PPARγ agonists in the EAE model (30), but is consistent with GW9662 displaying antiinflammatory properties in in vitro immune cell cultures and an LPS-triggered acute inflammation mouse model (31). In line with changes in EAE disease severity, *Scd1*^*-/-*^ induced changes in CNS inflammation and accumulation of CD3^+^ T cells were no longer apparent in the presence of GW9662 (Supplemental Figure 8, E-G). Furthermore, PPARγ antagonism reduced the observed increase in peripheral Treg frequency in *Scd1*^*-/-*^ EAE animals (Figure 6H and Supplemental Figure 7D). These findings provide evidence that *Scd1* deficiency impacts EAE disease severity, neuroinflammation, and Treg differentiation through PPARγ under physiological conditions.

## Discussion

An imbalance between pathogenic and protective T cell subsets is considered to propel autoimmunity. Findings from this study indicate that the FA desaturase SCD1 tightly controls T cell fate and autoimmunity. We report that pharmacological inhibition and genetic deficiency of SCD1 stimulates Treg differentiation, and, in parallel, attenuates neuroinflammation and disease severity in the EAE model. We further provide evidence that *Scd1* deficiency promotes Treg differentiation through ATGL-dependent hydrolysis of TGs and PC, thereby releasing non-esterified bioactive DHA, a natural ligand of the nuclear receptor PPARγ. Thus, our findings identify an intricate relationship between FA desaturation, Treg differentiation, and autoimmunity.

FA metabolism is increasingly being acknowledged as a hub and driver of T cell physiology in health and disease (4, 11-18). Our data indicate that the FA desaturase SCD1 acts as an endogenous brake on the differentiation of Tregs, thereby contributing to neuroinflammation in the EAE model. Consistent with our findings, previous studies showed that intracellular levels of substrates and products of SCD1, *i*.*e*. oleic (C18:1) and palmitic acid (C16:0), closely regulate T cell and Treg differentiation and maintenance (9, 13, 18). Likewise, SCD1 inhibition was recently found to increase the frequency of follicular Tregs in the spleen following influenza immunization (32). In contrast, *Scd1* deficiency accelerates and exacerbates the development of colitis by promoting the colitogenic potential of effector T cells (33). Here, elevated levels of SFA increased cellular membrane fluidity and augmented the secretory profile of effector T cells. However, the authors did not assess the impact of *Scd1* deficiency on Treg physiology. Differences in the immunopathology and applied experimental animal models, *i*.*e*. adoptive transfer of naïve or antigen-sensitized T cells to *Rag1*^-/-^ and wt mice, respectively, might well explain the opposing outcome of *Scd1* deficiency on disease initiation and severity in the colitis and EAE model. In summary, our findings highlight the importance of fatty acid desaturation by SCD1 in controlling Treg differentiation and protective autoimmunity in the EAE model.

Our findings demonstrate that lack of SCD1 increases ATGL-dependent hydrolysis of TGs and PC, resulting in elevated intracellular levels of non-esterified DHA, activation of PPARγ, enhanced differentiation of Tregs, and reduced EAE disease severity. Accordingly, ATGL is reported to favor the hydrolysis of TGs and phospholipids rich in PUFAs such as DHA, and DHA is a potent endogenous ligand for PPARγ (26, 34). Furthermore, previous studies showed that DHA and synthetic PPARγ agonists promote the differentiation and maintenance of Tregs, and reduce neuroinflammation in experimental animal models (29, 35-40). Surprisingly, we find that PPARγ antagonism reduces EAE disease severity as compared to control mice. The latter finding contradicts with the beneficial impact of PPARγ agonists in the EAE model but is consistent with GW9662 displaying anti-inflammatory properties in vitro and in vivo (31). All in all, our findings identify the SCD1-ATGL-PPAR axis as a key signaling pathway in driving Treg differentiation and highlight the complexity of PPAR signaling in (auto)immune disorders.

The molecular mechanisms that underlie increased ATGL-dependent release of DHA in *Scd1*^*-/-*^ T cells remain to be clarified. In the absence of SCD1, T cells may increase ATGL-dependent intracellular release of free DHA to protect themselves against endoplasmic reticulum stress induced by elevated levels of SFAs, potentially through activation of PPARγ (41, 42). In support of this notion, pathway analysis showed that differentially expressed genes in *Scd1*^*-/-*^ T cells were enriched in the canonical pathway ‘unfolded protein response’, a cellular response associated with ER stress (43). Alternatively, or in parallel, changes in the activity or abundance of signal transducer and activator of transcription 5 (STAT5), SP1, insulin, and TNFα may increase ATGL expression and activity in the absence of SCD1 (44-47). On a related note, while SFAs are generally regarded to promote inflammation, our findings implicate the opposite. Lack of SCD1 enhanced Treg differentiation and suppressed neuroinflammation, despite decreasing the intracellular desaturation index. Given that PUFAs are more potent PPARγ agonists than MUFAs (34), elevated DHA levels may outweigh and nullify the inflammatory impact of SFAs in our study. Alternatively, while the majority of studies determined the effects of exogenous SFAs on T cell physiology, the mode of action and cellular outcome of exogenously and endogenously altered SFAs levels might differ (48). Collectively, our findings indicate that the ATGL-DHA-PPARγ signaling axis is essential in driving enhanced Treg differentiation in *Scd1*-deficient T cells. However, more research is warranted to define the molecular mechanisms that control the activity of the ATGL-DHA-PPARγ axis.

LC-ESI-MS/MS analysis demonstrated that lack of SCD1 decreases the bulk of major fatty-acid containing lipid species in T cells, without affecting cell size and granularity. Given that differentially expressed genes in *Scd1*^-/-^ naïve T cells were enriched in pathways related to lipid efflux, these findings may reflect an increased capacity of *Scd1*^-/-^ T cells to dispose intracellular cholesterol and phospholipids. A complementary reduction in lipogenesis and activation in lipolysis may contribute to the observed decreased lipid load (49). Alongside elevated intracellular levels of non-esterified DHA, LC-MS/MS analysis revealed an increased abundance of the ω6 PUFA AA (C20:4) and its derivatives 11-HETE and 15-HETE in *Scd1*^-/-^ naïve T cells. Given that ATGL preferentially hydrolyses lipid species rich in PUFAs such as DHA and AA, these findings are consistent with increased ATGL activity in *Scd1*^-/-^ naïve T cells. Of interest, previous studies demonstrated that AA and its derivatives are endogenous PPARγ agonists and closely associated with Treg differentiation (34, 50, 51). To what extent AA, 11-HETE, and 15-HETE contribute to enhanced Treg differentiation in *Scd1*^-/-^ T cells remains to be determined.

A growing body of epidemiologic, preclinical, and observational studies suggest that obesity and Western diets rich in saturated fats, cholesterol, and carbohydrates are associated with Treg dysfunction, neuroinflammation, and autoimmunity (5, 6, 52-54). Of interest, obesity and consumption of a Western diet result in markedly elevated expression and activity of hepatic SCD1 (55-57), and *Scd1*-deficient mice are protected from diet-induced obesity (58, 59). Hence, our findings argue for SCD1 being an enzymatic driver underlying the impact of obesity and Western-type diets on immune cell imbalances and disease progression in autoimmune disorders such as MS. Vice versa, given that DHA inhibits *Scd1* expression (60), the beneficial impact of DHA supplementation on Treg differentiation, EAE severity, and disease pathology of autoimmune disorders such as MS, systemic lupus erythematosus, and rheumatoid arthritis might depend on changes in SCD1 activity (37, 61). Future dietary studies are warranted to unravel the causal role of SCD1 in driving the beneficial and detrimental impact of DHA and a Western diet, respectively, on the immunopathology of MS and other autoimmune disorders.

In summary, we report that SCD1 plays a crucial role in autoimmunity by suppressing Treg differentiation. Conclusions from this study extend our previous findings where we demonstrated that SCD1 impairs the reparative properties of macrophages and microglia in the CNS (20). Together, our findings argue for pharmacological inhibitors of SCD1 being a promising therapeutic strategy to simultaneously suppress autoimmunity, reduce neuroinflammation, and promote CNS repair. Our findings further provide a potential molecular rationale for the beneficial and detrimental impact of dietary factors on autoimmunity, and validate other recent studies that identified changes in fatty acid metabolism as a crucial determinant of the pathological outcome of these factors. Finally, this study endorses the essential role of PPARγ in driving autoimmunity and disease progression in MS (62-66). Altogether, our findings place SCD1 at the crossroad of autoimmunity and lipid metabolism.

## Materials and methods

### Mice

*Scd1*-deficient (*Scd1*^-/-^) mice and mice having the third exon of the *Scd1* gene flanked by *lox*P sites (*Scd1*^fl+/+^) are described in previous studies (67, 68). Both mouse strains were backcrossed at least 10 times with C57BL/6J mice. To generate phagocyte-specific *Scd1*^*-/-*^ mice, *Scd1*^fl+/+^ mice were crossbred with C57BL/6J *LysM*^Cre^ mice, which were kindly provided by prof. dr. Geert van Loo (VIB-UGent Center for Inflammation Research, University of Ghent, Belgium) (69). All mice were genotyped by PCR, as previously described (68), or according to protocols established by Jackson Laboratories. In all experiments using knockout mice, wild-type (wt) littermates were used as control. For the experiment using phagocyte-specific knockout mice, *LysM*^Cre+/-^ and *Scd1*^fl+/+^ mice were used as control. For the adoptive transfer experiments, wt C57BL/6J animals were purchased from Envigo. Mice were maintained on a 12 h light/dark cycle with free access to water and a standard chow diet.

### Experimental autoimmune encephalomyelitis (EAE)

At the age of 9-12 weeks, female mice were immunized subcutaneously with myelin oligodendrocyte glycoprotein peptide 35-55 (MOG^35-55^) emulsified in complete Freund’s adjuvant supplemented with *Mycobacterium tuberculosis* according to manufacturer’s guidelines (EK-2110 kit; Hooke Laboratories). Directly after immunization and after 24 h, mice were intraperitoneally (i.p.) injected with 100 ng or 50 ng pertussis toxin (depending on lot number). Starting 5 days post-immunization, EAE mice were treated daily with an SCD1 inhibitor (CAY10566, 2.5 mg/kg, every 12 hours by oral gavage), ATGL inhibitor (Atglistatin, 200 µmol/kg, every 12 hours by oral gavage), PPARγ inhibitor (GW9662, 2mg/kg, every 24 hours by i.p. injection), or vehicle (PBS for i.p., methylcellulose for oral gavage). Mice were weighed and clinically evaluated for neurological signs of the disease on a daily basis following a five-point standardized rating of clinical symptoms (0: no clinical symptoms; 1: tail paralysis; 2: tail paralysis and partial hind limb paralysis; 3: complete hind limb paralysis; 4: paralysis to the diaphragm; 5: death by EAE). For the adoptive transfer experiments, inguinal lymph nodes were isolated from EAE donor mice at day 10 post-immunization. Single-cell suspensions were obtained by pushing the tissue through a 70 µm cell strainer and subsequently cultured at 7 × 10^6^ cells/ml in stimulation medium consisting of RPMI1640 supplemented with 0.5% penicillin/streptomycin (Gibco), 10% FCS (Gibco), 1% non-essential amino acid (NEAA, Sigma-Aldrich), 1% sodium pyruvate (Sigma-Aldrich), 20 ng/ml IL23 (BioLegend), and 20 µg/ml rMOG_35-55_ (Hooke Laboratories). After three days, 10 × 10^6^ cells were intraperitoneally injected into wt or *Scd1*^-/-^ recipient mice, which were weighed and clinically evaluated for neurological signs of the disease on a daily basis.

### Mouse T cell cultures

CD4^+^ T cells were isolated *ex vivo* from spleens of 9-12-week-old wt and *Scd1*^-/-^ mice, using the CD4^+^ T Cell Isolation Kit (Miltenyi Biotec) according to manufacturer’s instructions. Next, live naïve (CD4^+^CD25^-^CD44^med^CD62L^+^) T cells were obtained using a FACSAria cell sorter (BD Biosciences). The purity of the isolated cells was > 95%. For Treg induction, 0.75 × 10^5^ naïve T cells were cultured for one or two days with plate-bound anti-CD3ε (2 µg/ml, clone 145-2C11; BD Biosciences) in differentiation medium consisting of RPMI1640 supplemented with 1% penicillin/streptomycin, 10% FCS, 1% NEAA, 1% L-glutamine (Sigma-Aldrich), 50 µM 2-mercaptoethanol (Gibco), anti-CD28 (2 µg/ml, clone 37.51; BD Biosciences), and rmTGFβ (10 ng/ml; R&D Systems) in a 96-well U-bottom plate. SCD1 inhibitor (CAY10566, 1 µM; Cayman Chemicals), ATGL inhibitor (Atglistatin, 20 µM, synthesized as described previously (70)), non-esterified DHA (C22:6, 1 µM; Cayman Chemicals), PPARα antagonist (GW6471, 25 µM; Sigma-Aldrich), PPARβ antagonist (PT-S58, 25 µM; Sigma-Aldrich), PPARγ antagonist (GW9662, 25 µM; Sigma-Aldrich) or vehicle were added daily starting from the onset of cultures. DHA was dissolved in ethanol and complexed to fatty acid-free BSA (Sigma-Aldrich) in a 4:1 molar ratio. To define the lipidome after ATGL inhibition, 0.75 × 10^5^ *Scd1*^-/-^ naïve T cells were cultured for two days in RPMI1640 supplemented with 1% penicillin/streptomycin, 10% FCS, 1% NEAA, 1% L-glutamine (Sigma-Aldrich), 50 µM 2-mercaptoethanol (Gibco), and rmIL7 (10 ng/ml, Peprotech) in a 96-well U-bottom plate. ATGL inhibitor or vehicle were added daily starting from the onset of cultures.

Suppressive capacity of wt and *Scd1*^-/-^ Tregs was assessed as described previously (71). In brief, antigen presenting cells (APCs, CD4^-^), effector T cells (Teff, CD4^+^CD25^-^), and Tregs (CD4^+^CD25^+^) were isolated *ex vivo* from spleens of 11-week-old wt and *Scd1*^-/-^ animals, using the CD4^+^CD25^+^ Regulatory T Cell Isolation Kit (Miltenyi Biotec) according to manufacturer’s instructions. The purity of the isolated cells was > 95%. Increasing amounts of Tregs were co-cultured alongside 0.5 × 10^5^ wt CFSE-labeled Teff and 2 × 10^5^ irradiated wt APCs (30 Gy) in a 96-well V-bottom plate for three days. RPMI1640 supplemented with 1% penicillin/streptomycin, 10% FCS, 1% NEAA, 1% L-glutamine, and 50 µM 2-mercaptoethanol was used as culture medium. To characterize the T cell compartment of wt and *Scd1*^-/-^ EAE animals, spleen and inguinal lymph nodes were isolated from EAE mice at day 10 post-immunization. Single-cell suspensions were obtained by pushing the tissue through a 70 µm cell strainer and cultured overnight at 5 × 10^6^ cells/ml in culture medium (RPMI1640 supplemented with 0.5% penicillin/streptomycin, 10% FCS, 1% NEAA, and 1% sodium pyruvate).

### Human T cell cultures

Using density gradient centrifugation, peripheral blood mononuclear cells (PBMCs) were obtained from healthy donors. CD4^+^CD25^-^ T cells were isolated using the CD4^+^ T Cell Isolation Kit (Miltenyi Biotec) and CD25 Microbeads (Miltenyi Biotec) according to manufacturer’s instructions. Next, recent thymic emigrated (CD4^+^CD127^+^CD31^+^, designated hereafter as naïve) T cells were obtained using a FACSAria cell sorter (BD Biosciences). The purity of the isolated cells was > 95%. For Treg induction, 0.1 × 10^6^ naïve T cells were cultured for four days in differentiation medium consisting of RPMI1640 supplemented with 0.5% penicillin/streptomycin, 10% FCS, 1% NEAA, 1% sodium pyruvate, IL2 (50 U/ml, Roche), rhTGFβ (5 ng/ml, Peprotech), soluble anti-CD3ε (1.25 µg/ml, clone OKT3, eBioscience), and Treg Suppression Inspector beads (Miltenyi Biotec) in a 96-well U-bottom plate. SCD1 inhibitor (CAY10566, 1 µM), ATGL inhibitor (Atglistatin, 20 µM, synthesized as described previously (70)), PPARα antagonist (GW6471, 25 µM; Sigma-Aldrich), PPARβ antagonist (PT-S58, 25 µM; Sigma-Aldrich), PPARγ antagonist (GW9662, 10 µM), or vehicle were added daily starting from the onset of cultures.

Suppressive capacity of Tregs was assessed as described previously (72). In brief, effector T cells (Teff, CD4^+^CD25^-^) were stained with CellTrace Violet (CTV, ThermoFisher Scientific, 2.5 µM) and cultured with Tregs (CD4^+^CD25^+^) which were differentiated in the presence of SCD1 inhibitor or vehicle. T cells were isolated using the CD4^+^ T Cell Isolation Kit (Miltenyi Biotec) and CD25 Microbeads (Miltenyi Biotec) according to manufacturer’s instructions. The purity of the isolated cells was > 95%. Increasing amounts of Tregs were co-cultured alongside 0.5 × 10^5^ effector T cells in a 96-well U-bottom plate for 5 days before being analysed by FACS. RPMI1640 supplemented with 0.5% penicillin/streptomycin, 10% FCS, 1% NEAA, 1% sodium pyruvate, IL2, rhTGFβ, and Treg Suppression Inspector beads was used as culture medium.

### shRNA mediated gene silencing

Lentiviral particles were produced in HEK293T cells by co-transfecting packaging vector psPAX2 (8µg, Addgene, #12260), envelop vector pMD2.G (4µg, Addgene, #12259), and lentiviral vector (12µg, Mission Sigma) expressing the indicated specific shRNA or a control vector expressing an unspecific shRNA, using 125 mM CaCl_2_. The specific shRNA clones were TRCN0000249774 for ATGL, or TRC0000001657 for PPARγ. Supernatants were collected over the course of three days and filtered through a 0.45 μm filter. Lentiviral particles were concentrated by ultracentrifugation for 2 h at 70,000 x g. 0.75 × 10^5^ wild-type and *Scd1*^-/-^ naïve T cells/well were stimulated for one day prior infection. Next, T cells were transduced with lentiviral particles at a lentiviral particle medium:culture medium ratio of 1:6. One day post-infection, naïve T cells were stimulated with rmTGFβ for Treg induction. Knockdown of endogenous ATGL and PPARγ was confirmed using flow cytometry.

### Cell lines

Jurkat T cells were cultured in RPM1640 supplemented with 0.5% penicillin/streptomycin, 10% FCS, and 1% L-glutamine.

### Flow cytometry

For the staining of spleen- and lymph node-derived lymphocytes, the following antibodies were used (all purchased from BioLegend): CD45 Alexa Fluor 700 (clone 30-F11, 1:200), CD3 FITC (clone 17A2, 1:200), CD4 Pacific Blue (clone GK1.5, 1:200), CD8 BV510 (clone 53-6.7, 1:200), IFNγ PE-Cy7 (clone XMG1.2, 1:200), IL17 PE-Dazzle 594 (clone TC1-18H10.1, 1:200), IL4 PE (clone 11B11, 1:50), and FOXP3 Alexa Fluor 647 (clone MF-14, 1:200). For the isolation of mouse naïve T cells and the subsequent staining of differentiated Tregs, the following antibodies were used: CD4 PerCP-Cy5.5 (clone RM4-5, 1:100; BioLegend), CD44 APC-Cy7 (clone IM7, 1:100; BioLegend), CD25 PE-Cy7 (clone PC61, 1:200; BD Biosciences), CD62L APC (clone MEL1, 1:100; BioLegend), and FOXP3 eFluor450 (clone FJK-16s, 1:30, Invitrogen). To assess murine Treg suppressive capacity, the following antibodies were used: CD4 Pacific Blue (clone GK1.5, 1:200; BioLegend), CD25 PE-Cy7 (clone PC61, 1:200; BD Biosciences), FOXP3 Alexa Fluor 647 (clone MF-14, 1:200; BioLegend). To assess ATGL and PPARγ abundance, the following antibodies were used: ATGL (NB110-41536, 1:100, Novus), PPARγ (sc-7273, 1:100, Santa Cruz Biotechnology). For the isolation of human naïve T cells and the subsequent staining of differentiated Tregs, the following antibodies were used: CD4 APC-eFluor780 (clone OKT4, 1:20; eBioscience), CD31 Alexa Fluor 488 (clone WM59, 1:20; BioLegend), CD127 PE (eBioRDR5, 1:20; Invitrogen), CD25 PE-Cy7 (clone M-A251, 1:40; BioLegend), FOXP3 BV421 (clone 206D, 1:40; BioLegend). Dead cells were excluded by incubation with Propidium Iodide (1:250, BD Biosciences), Fixable Viability Dye eFluor506 (1:1000, ThermoFisher Scientific), or Fixable Viability Dye Zombie NIR (1:1000, BioLegend). To analyze surface markers, cells were stained in 1x PBS containing 2% FCS and 0.1% azide. To stain for intracellular cytokines, cells were stimulated with phorbol 12-myristate 13-acetate (20 ng/ml; Sigma-Aldrich), Calcium Ionomycin (1 µg/ml; Sigma-Aldrich), and Golgiplug (2 µg/ml; BD Biosciences) for 4 h and the FOXP3 / Transcription Factor staining buffer set (Invitrogen) was used according to manufacturer’s instructions. Cells were acquired on an LSRFortessa Flow Cytometer (BD Biosciences).

### Quantitative PCR (qPCR)

Tissue or cells were lysed using QIAzol (Qiagen). RNA extraction and synthesis of complementary DNA was performed as described previously (20). qPCR was subsequently conducted on a StepOnePlus or QuantStudio 3 detection system (Applied Biosystems). Data was analyzed using the ΔΔCt method and normalized to the most stable reference genes, as described previously (73). Primer sequences are available on request.

### Immunofluorescence microscopy and image analysis

Cryosections were fixed in acetone for 10 min. Immunostaining and analysis of cryosections were performed as described previously (74). Immune cell infiltrates were stained using the following antibodies: anti-CD3 (1:150; Bio-Rad) and anti-F4/80 (1:100; Bio-Rad), combined with the secondary Alexa Fluor 555-labeled anti-rat IgG antibody (1:400; Invitrogen). Analysis was carried out using a Nikon eclipse 80i microscope and ImageJ software.

### Bulk RNA sequencing

Tissue or cells were lysed using QIAzol. RNA was extracted using the RNeasy mini kit (Qiagen). RNA samples were processed by the Genomics Core Leuven (Belgium). Total RNA content was analyzed with Nanodrop 1000 spectrophotometer (ThermoFisher Scientific) and RNA integrity was evaluated using Bioanalyzer (Agilent). Sequencing libraries were generated using the Lexogen’s QuantSeq kit and sequenced on the Illumina HiSeq4000 sequencing system, generating 50 bp single-end reads. Splice-aware alignment was performed with STAR (v2.6.1b) using the default parameters (75). Reads mapping to multiple loci in the reference genome are discarded. Quantification of reads per gene was performed with HT-seq Count v2.7.14. For count-based differential expression analysis of a single gene, R-based Bioconductor package DESeq2 (The R Foundation for Statistical Computing) was used. Data are available as a GEO dataset under the accession number GSE160040. Differentially expressed genes were identified based on a log_2_ fold change < -0.5 and > 0.5, and a P value < 0.05. The list of differentially expressed genes was used as the input for the analysis with QIAGEN’s Ingenuity Pathway Analysis (IPA, Qiagen).

### Liquid chromatography electrospray ionization tandem mass spectrometry (LC-ESI-MS/MS)

Naïve T cell pellets were reconstituted in 700 μl 1x PBS, mixed with 800 μl 1 N hydrochloric acid:methanol (MeOH) 1:8 (v/v), 900 μl chloroform, 200 μg/ml of the antioxidant 2,6-di-tert-butyl-4-methylphenol (BHT; Sigma-Aldrich), and 3 μl of SPLASH LIPIDOMIX Mass Spec Standard (#330707, Avanti Polar Lipids). After vortexing and centrifugation, the lower organic fraction was collected and evaporated using a Savant Speedvac spd111v (Thermo Fisher Scientific) at room temperature and the remaining lipid pellet was stored at - 20°C under argon. The lipid pellet was reconstituted in 100% ethanol. Lipid species were analyzed by LC-ESI-MS/MS on a Nexera X2 UHPLC system (Shimadzu) coupled with hybrid triple quadrupole/linear ion trap mass spectrometer (6500+ QTRAP system; AB SCIEX). Chromatographic separation was performed on a XBridge amide column (150 mm × 4.6 mm, 3.5 μm; Waters) maintained at 35°C using mobile phase A [1 mM ammonium acetate in H_2_O-acetonitrile 5:95 (v/v)] and mobile phase B [1 mM ammonium acetate in H_2_O-acetonitrile 50:50 (v/v)] in the following gradient: (0-6 min: 0% B ➡ 6% B; 6-10 min: 6% B ➡ 25% B; 10-11 min: 25% B ➡ 98% B; 11-13 min: 98% B ➡ 100% B; 13-19 min: 100% B; 19-24 min: 0% B) at a flow rate of 0.7 ml/min which was increased to 1.5 ml/min from 13 min onwards. Sphingomyelin (SM), cholesteryl esters (CE), ceramides (CER), dihydroceramides (DCER), hexosylceramides (HCER), lactosylceramides (LCER) were measured in positive ion mode with a precursor scan of 184.1, 369.4, 264.4, 266.4, 264.4, and 264.4 respectively. Tri-, di-, and mono-acylglycerides (TGs, DGs, and MGs) were measured in positive ion mode with a neutral loss scan for one of the fatty acyl moieties. Phosphatidylcholine (PC), lysophosphatidylcholine (LPC), phosphatidylethanolamine (PE), lysophosphatidylethanolamine (LPE), phosphatidylglycerol (PG), phosphatidylinositol (PI), and phosphatidylserine (PS) were measured in negative ion mode by fatty acyl fragment ions. Lipid quantification was performed by scheduled multiple reactions monitoring (MRM), the transitions being based on the neutral losses or the typical product ions as described above. The instrument parameters were as follows: Curtain Gas = 35 psi; Collision Gas = 8 a.u. (medium); IonSpray Voltage = 5500 V and −4,500 V; Temperature = 550°C; Ion Source Gas 1 = 50 psi; Ion Source Gas 2 = 60 psi; Declustering Potential = 60 V and −80 V; Entrance Potential = 10 V and −10 V; Collision Cell Exit Potential = 15 V and −15 V. Peak integration was performed with the MultiQuantTM software version 3.0.3. Lipid species signals were corrected for isotopic contributions (calculated with Python Molmass 2019.1.1) and were quantified based on internal standard signals and adheres to the guidelines of the Lipidomics Standards Initiative (LSI; level 2 type quantification as defined by the LSI). Only the detectable lipid classes and fatty acyl moieties are reported in this manuscript.

### Liquid chromatography tandem mass spectrometry (LC-MS/MS)

Protein precipitation was performed on naïve T cells by dissolving them in 500 µl MeOH, to which 4 µl of internal standard solution consisting of [^2^H_4_]LTB_4_, [^2^H_8_]15-HETE, [^2^H_4_]PGE_2_, and [^2^H_5_]DHA (50 ng/ml) was added. Samples were placed in -20°C for 20 min to equilibrate after which they were spun down for 10 min (16200 g at 4°C). Supernatant was diluted with 2.5 ml H_2_O and the pH was adjusted to 3.5 using formic acid (99%). Lipids were extracted using solid-phase extraction (SPE). In short, C18 cartridges (Sep-Pak, Vac3 3cc (200mg)) were equilibrated with MeOH (LC-MS grade, Merck) and H_2_O (LC-MS grade, VWR Chemicals), after which the samples were loaded. Subsequently, cartridges were washed with H_2_O (LC-MS grade) and *n*-hexane (Sigma-Aldrich), followed by elution using methylformate (spectrophotometric grade, Sigma-Aldrich). The elutes were then dried at 40°C using a gentle stream of nitrogen and reconstituted in 100 µl MeOH:H_2_O (40%). Next, samples were analyzed using a targeted LC-MS/MS method as described previously (76). Here, chromatographic separation was accomplished using a Shimadzu LC-system, consisting of two LC-30AD pumps, a SIL-30AC autosampler, and a CTO-20AC column oven (Shimadzu). Compounds were separated on a Kinetex C18 column (50 mm × 2.1 mm, 1.7 µm), protected with a C8 pre-column (Phenomenex). Samples were eluted at a constant flowrate of 400 µl/min with a gradient of H_2_O (eluent A) and MeOH (eluent B), both with 0.01% acetic acid (76). Other instrument parameters were in line with the protocol of Jónasdóttir et al. (76) and scheduled MRM mode was used to detect compounds. Compounds were identified using their relative retention times together with characteristic mass transitions. These and other individually optimized parameters can be found in Supplementary Table 1. Only the detectable non-esterified fatty acids and downstream metabolites were reported in this manuscript.

### Luciferase-based nuclear receptor reporter assay

To determine ligation of PPARα, PPARβ, and PPARγ, luciferase-based reporter assays were performed using the ONE-GloTM Luciferase Assay System kit (Promega). Jurkat T cells were transfected with bacterial plasmid constructs expressing luciferase under control of the promotor region of the ligand-binding domain for PPARα, PPARβ, or PPARγ, which were kindly provided by prof. dr. Bart Staels (University of Lille, Inserm, France) (77, 78). Cells were grown to 50–60% confluency in 60 mm plates, transfected with 1.8 μg of plasmid DNA including 0.2 μg pGAL4hPPARα, pGAL4hPPARβ, or pGAL4hPPARγ, 1 μg pG5-TK-GL3, and 0.6 μg of pCMV-β-galactosidase, using JetPEI (Polyplus-transfection SA, France) as transfection reagent. Transfected cells were treated with SCD1 inhibitor (CAY10566, 10 µM), ATGL inhibitor (Atglistatin, 20 µM), non-esterified DHA (1 µM), PPARα agonist (WY-14643, 10 µM; Sigma-Aldrich), PPARβ agonist (GW501516, 10 µM; Sigma-Aldrich), PPARγ agonist (Rosiglitazone, 10 µM; Millipore), or vehicle for 24 h. Following treatment, cells were lysed in lysis buffer (25 mM Glycyl-Glycine, 15 mM MgSO4, 4 mM EGTA, and 1x Triton; all from Sigma-Aldrich). To correct for transfection efficacy, β-galactosidase activity was measured using lysate diluted 1:10 in B-gal buffer, consisting of 20% 2-Nitrophenyl β-D-galactopyranoside (ONGP; Sigma-Aldrich) and 80% Buffer Z (0.1 M Na2HPO4, 10 mM KCl, 1 mM MgSO4, and 3.4 µl/ml 2-mercaptoethanol; all from Sigma-Aldrich). Luminescence and absorbance (410 nm) were measured using the FLUOstar Optima (BMG Labtech).

### Statistics

Data was analyzed for statistical significance using GraphPad Prism and are reported as mean ± SEM. Data were tested for normal distribution using the d’Agostino and Pearson omnibus normality test. When data sets were normally distributed, an ANOVA (Tukey’s post hoc analysis) or a two-tailed unpaired student T-test (with Welch’s correction if necessary) was used to determine statistical significance between groups. If data sets were not normally distributed, the Kruskal-Wallis or Mann-Whitney analysis was used. Significant differences were identified by P values < 0.05 (*P < 0.05, **P < 0.01, and ***P < 0.001).

### Study approval

All animal procedures were conducted in accordance with the institutional guidelines and approved by the Ethical Committee for Animal Experiments of Hasselt University (ID201521, ID201611, ID201822, ID201828, ID201912K, ID201914). All experimental protocols using PBMCs were conducted in accordance with institutional guidelines and approved by the Medical Ethical Committee of Hasselt University (UH-IMMVET-P1). Written informed consents were obtained from all participants included in this study.

## Author contributions

EG, ML, MH, MK, JJAH, and JFJB conceived experiments. EG, PB, BFC-R, ML, JD, JYB, LVZ, and JFJB performed experiments. EG, PB, ML, IH, MH, JD, JYB, and JFJB analyzed data. EG, PB, BFC-R, ML, IH, MH, JD, JYB, MG, LVZ, JMN, NH, GK, RZ, RB, JVS, BB, MK, JJAH, and JFJB discussed results. JMN, PS, NH, MG, RZ, RB, GK, and MK contributed reagents, materials, and analysis tools. EG, MH, JJAH, and JFJB wrote the manuscript. EG, PB, BFC-R, ML, IH, MH, JD, JYB, LVZ, JMN, PS, NH, MG, RZ, RB, GK, JVS, BB, MK, JJAH, and JFJB revised the manuscript.

## Acknowledgements

We thank MP Tulleners for excellent technical assistance. We also like to thank other members in the Biomedical Research Institute (Hasselt University) for providing feedback and suggestions during preparation of the manuscript.

