## Supplemental Tables and Figures for "Fatty acid desaturation by stearoyl-CoA desaturase-1 controls regulatory T cell differentiation and autoimmunity"

### Supplementary Table 1: LC-MS/MS parameters

| Group | Compound | Lipid Maps ID | Retention time [min] | m/z in Q1 | m/z in Q3 | Declustering potential [V] | Collision energy [V] | Collision cell exit potential [V] |
|---|---|---|---|---|---|---|---|---|
| Epoxyeicosatrienoic acids (EET) | 14(15)-EET | LMFA03080005 | 8.1 | 319 | 218.9 | -5 | -16 | -55 |
| Hydroxydocosahexaenoic acids (HDHA) | 14(S)-HDHA | LMFA04000058 | 8 | 343.1 | 204.9 | -60 | -18 | -27 |
| Hydroxydocosahexaenoic acids (HDHA) | 17-HDHA | LMFA04000072 | 7.9 | 343.1 | 245 | -65 | -16 | -15 |
| Hydroxyeicosapentaenoic acids (HEPE) | 12-HEPE | LMFA03070031 | 7.6 | 317 | 179 | -60 | -18 | -17 |
| Hydroxyeicosapentaenoic acids (HEPE) | 15-HEPE | LMFA03070009 | 7.5 | 317.1 | 219 | -65 | -18 | -19 |
| Hydroxyeicosatetraenoic acids (HETE) | 11-HETE | LMFA03060003 | 7.9 | 319.1 | 167 | -70 | -22 | -15 |
| Hydroxyeicosatetraenoic acids (HETE) | 12-HETE | LMFA03060007 | 7.9 | 319.1 | 179 | -65 | -20 | -23 |
| Hydroxyeicosatetraenoic acids (HETE) | 15-HETE | LMFA03060001 | 7.8 | 319.1 | 219.1 | -55 | -18 | -9 |
| Hydroxyeicosatetraenoic acids (HETE) | 17-OH-DH-HETE | N/A | 8.2 | 347.1 | 247 | -110 | -22 | -27 |
| Hydroxyeicosatetraenoic acids (HETE) | 5-HETE | LMFA03060002 | 8 | 319.1 | 115 | -65 | -18 | -11 |
| Hydroxyeicosatetraenoic acids (HETE) | 8-HETE | LMFA03060006 | 7.9 | 319.1 | 154.9 | -70 | -20 | -19 |
| Hydroxyoctadecadienoic acids (HoDE) | 13-HoDE | LMFA02000228 | 7.7 | 295 | 194.9 | -110 | -24 | -21 |
| Hydroxyoctadecadienoic acids (HoDE) | 9-HoDE | LMFA02000188 | 7.7 | 295 | 171 | -130 | -22 | -7 |
| Hydroxyoctadecatrienoic acids (HoTrE) | 9-HoTrE | LMFA02000024 | 7.4 | 293 | 170.9 | -75 | -20 | -15 |
| Internal standards | 15-HETE-d8 | LMFA03060080 | 7.8 | 327.2 | 226 | -85 | -18 | -11 |
| Internal standards | DHA-d5 | LMFA01030762 | 8.8 | 332 | 288.1 | -75 | -16 | -13 |
| Internal standards | LTB4-d4 | LMFA03020030 | 6.9 | 339.1 | 196.9 | -70 | -22 | -19 |
| Internal standards | PGE2-d4 | LMFA03010008 | 4.9 | 355.1 | 193 | -50 | -26 | -17 |
| Keto-eicosatetraenoic acids (KETE/OxoETE) | 12-KETE | LMFA03060019 | 7.9 | 317 | 153 | -60 | -22 | -9 |
| Keto-eicosatetraenoic acids (KETE/OxoETE) | 15-KETE | LMFA03060051 | 7.8 | 317 | 113 | -10 | -22 | -5 |
| Keto-eicosatetraenoic acids (KETE/OxoETE) | 5-KETE | LMFA03060011 | 8.1 | 317 | 203.1 | -70 | -24 | -11 |
| Leukotrienes (LT) | 6-trans-LTB4 | LMFA03020013 | 6.7 | 335.1 | 194.9 | -105 | -22 | -11 |
| Polyunsaturated fatty acids | AA | LMFA01030001 | 8.8 | 303 | 205.1 | -155 | -20 | -11 |
| Polyunsaturated fatty acids | AdA | LMFA01030178 | 9.1 | 331.1 | 233 | -130 | -22 | -11 |
| Polyunsaturated fatty acids | ALA/GLA | LMFA01030152 / LMFA01030141 | 8.6 | 277 | 233 | -90 | -22 | -29 |
| Polyunsaturated fatty acids | DGLA | LMFA01030158 | 9 | 305.1 | 261.2 | -85 | -22 | -13 |
| Polyunsaturated fatty acids | DHA | LMFA01030185 | 8.8 | 327.1 | 229.2 | -115 | -18 | -11 |

| | | | | | | | | |
|---|---|---|---|---|---|---|---|---|
| Polyunsaturated fatty acids | DPAn-3 | LMFA04000044 | 8.9 | 329.1 | 231.1 | -50 | -20 | -17 |
| Polyunsaturated fatty acids | DPAn-6 | LMFA01030182 | 9 | 329.1 | 231.1 | -50 | -20 | -17 |
| Polyunsaturated fatty acids | EPA | LMFA01030759 | 8.6 | 301 | 202.9 | -125 | -18 | -21 |
| Polyunsaturated fatty acids | LA | LMFA01030120 | 8.8 | 279 | 261 | -115 | -28 | -13 |
| Prostaglandins (PG) | 15-deoxy-PGJ2 | LMFA03010021 | 7.3 | 315 | 203 | -50 | -28 | -19 |
| Prostaglandins (PG) | 15-keto-PGE2 | LMFA03010030 | 4.5 | 349 | 234.9 | -65 | -20 | -13 |
| Prostaglandins (PG) | PGD2 | LMFA03010004 | 5 | 351.1 | 233 | -30 | -16 | -13 |
| Prostaglandins (PG) | PGE2 | LMFA03010003 | 4.9 | 351.2 | 271.1 | -50 | -22 | -21 |
| Prostaglandins (PG) | PGF2alpha | LMFA03010002 | 5.2 | 353.1 | 193 | -80 | -34 | -11 |
| Prostaglandins (PG) | PGJ2 | LMFA03010019 | 6.1 | 333 | 271 | -30 | -22 | -17 |
| Thromboxanes (Tx) | TXB2 | LMFA03030002 | 4.6 | 369.1 | 169 | -55 | -24 | -15 |

**Supplementary Table 2: Differentially expressed genes in *Scd1*-deficient CD4⁺ naïve T cells vs. wt CD4⁺ naïve T cells**

| Upregulated genes | | | | | |
|---|---|---|---|---|---|
| **Symbol** | **Entrez Gene Name** | **Basemean** | **Log2(Fold change)** | **P value** | **Padj** |
| Dapl1 | death associated protein like 1 | 402.509 | 0.589 | 0.00001 | 0.194 |
| Ctsw | cathepsin W | 27.174 | 1.363 | 0.00008 | 0.377 |
| Igha | immunoglobulin heavy constant alpha | 7.323 | 3.633 | 0.00018 | 0.65 |
| Klhdc2 | kelch domain containing 2 | 102.137 | 0.698 | 0.00032 | 0.766 |
| F2rl1 | F2R like trypsin receptor 1 | 49.369 | 0.855 | 0.00048 | 0.975 |
| Irf2bpl | interferon regulatory factor 2 binding protein like | 102.321 | 0.579 | 0.00100 | 0.977 |
| Trim30d | tripartite motif-containing 30A | 27.963 | 1.107 | 0.00106 | 0.977 |
| Bbs12 | Bardet-Biedl syndrome 12 | 3.15 | 4.564 | 0.00110 | 0.977 |
| Mir22hg | Mir22 host gene (non-protein coding) | 7.688 | 2.053 | 0.00206 | 1 |
| Ikzf2 | IKAROS family zinc finger 2 | 35.539 | 0.971 | 0.00240 | 1 |
| Foxp3 | forkhead box P3 | 9.53 | 1.789 | 0.00293 | 1 |
| Npc1 | NPC intracellular cholesterol transporter 1 | 46.302 | 0.816 | 0.00296 | 1 |
| Neurl3 | neuralized E3 ubiquitin protein ligase 3 | 55.838 | 0.753 | 0.00344 | 1 |
| Ppp6r3 | protein phosphatase 6 regulatory subunit 3 | 73.903 | 0.613 | 0.00347 | 1 |
| C330007P06Rik | chromosome X open reading frame 56 | 10.488 | 1.714 | 0.00439 | 1 |
| Abca1 | ATP binding cassette subfamily A member 1 | 16.024 | 1.296 | 0.00485 | 1 |
| Gab3 | GRB2 associated binding protein 3 | 35.96 | 0.804 | 0.00525 | 1 |
| Kif1b | kinesin family member 1B | 175.283 | 0.548 | 0.00541 | 1 |
| Ipo7 | importin 7 | 11.185 | 1.395 | 0.00609 | 1 |
| 1300002E11Rik | RIKEN cDNA 1300002E11 gene | 27.268 | 0.872 | 0.00842 | 1 |
| Plekha3 | pleckstrin homology domain containing A3 | 37.796 | 0.76 | 0.00848 | 1 |
| 1810026B05Rik | RIKEN cDNA 1810026B05 gene | 24.673 | 0.947 | 0.00883 | 1 |
| Nacc1 | nucleus accumbens associated 1 | 14.25 | 1.212 | 0.00907 | 1 |
| Fam169b | family with sequence similarity 169, member B | 71.956 | 0.529 | 0.00980 | 1 |
| Prg4 | proteoglycan 4 (megakaryocyte stimulating factor, articular superficial zone protein) | 6.309 | 1.889 | 0.00997 | 1 |
| Cetn2 | centrin 2 | 57.202 | 0.641 | 0.0105 | 1 |
| Scyl2 | SCY1 like pseudokinase 2 | 32.451 | 0.788 | 0.0108 | 1 |
| Skil | SKI like proto-oncogene | 95.086 | 0.558 | 0.0108 | 1 |
| Gm11427 | schlafen 4 pseudogene | 1.346 | 4.088 | 0.0116 | 1 |
| Eif4enif1 | eukaryotic translation initiation factor 4E nuclear import factor 1 | 46.751 | 0.708 | 0.0117 | 1 |
| Pnpla2 | patatin like phospholipase domain containing 2 | 36.933 | 0.739 | 0.0129 | 1 |
| Gm7079 | ribosomal protein S29 pseudogene | 147.45 | 0.519 | 0.0129 | 1 |
| Aim2 | absent in melanoma 2 | 43.081 | 0.649 | 0.0134 | 1 |
| Spock2 | SPARC (osteonectin), cwcv and kazal like domains proteoglycan 2 | 2.305 | 3.258 | 0.0136 | 1 |
| Gm5244 | ribosomal protein, large, P1 pseudogene | 55.087 | 0.596 | 0.0157 | 1 |
| Cd55 | CD55 molecule (Cromer blood group) | 52.746 | 0.578 | 0.0169 | 1 |
| Ar | androgen receptor | 24.281 | 0.907 | 0.0170 | 1 |
| Gm10335 | ribosomal protein L23A pseudogene | 56.443 | 0.558 | 0.0178 | 1 |
| Ano6 | anoctamin 6 | 76.375 | 0.543 | 0.0179 | 1 |
| Ttc39b | tetratricopeptide repeat domain 39B | 64 | 0.589 | 0.0181 | 1 |
| Ptdss1 | phosphatidylserine synthase 1 | 46.612 | 0.597 | 0.0183 | 1 |
| Pik3c2a | phosphatidylinositol-4-phosphate 3-kinase catalytic subunit type 2 alpha | 45.332 | 0.623 | 0.0185 | 1 |
| Brca2 | BRCA2 DNA repair associated | 8.47 | 1.493 | 0.0188 | 1 |
| Rpl26-ps4 | ribosomal protein L26, pseudogene 4 | 7.112 | 1.485 | 0.0192 | 1 |
| Arhgef12 | Rho guanine nucleotide exchange factor 12 | 13.597 | 1.096 | 0.0194 | 1 |
| Snx13 | sorting nexin 13 | 13.856 | 1.041 | 0.0201 | 1 |

| | | | | | |
|---|---|---|---|---|---|
| H2-Q6 | major histocompatibility complex, class I, A | 24.728 | 0.938 | 0.0202 | 1 |
| H2-T24 | histocompatibility 2, T region locus 24 | 21.443 | 0.887 | 0.0202 | 1 |
| Sesn3 | sestrin 3 | 57.459 | 0.566 | 0.0205 | 1 |
| Golga2 | golgin A2 | 55.379 | 0.542 | 0.0208 | 1 |
| Zfp667 | zinc finger protein 667 | 5.37 | 1.793 | 0.0209 | 1 |
| Tmed4 | transmembrane p24 trafficking protein 4 | 22.171 | 0.869 | 0.0211 | 1 |
| Zcchc10 | zinc finger CCHC-type containing 10 | 30.908 | 0.764 | 0.0217 | 1 |
| Ammecr1l | AMMECR1 like | 71.228 | 0.544 | 0.0218 | 1 |
| Dlgap5 | DLG associated protein 5 | 1.873 | 3.784 | 0.0220 | 1 |
| 1810044D09Rik | RIKEN cDNA 1810044D09 gene | 2.692 | 2.884 | 0.0221 | 1 |
| Usp37 | ubiquitin specific peptidase 37 | 27.807 | 0.743 | 0.0221 | 1 |
| Heca | hdc homolog, cell cycle regulator | 54.187 | 0.586 | 0.0226 | 1 |
| Klhl8 | kelch like family member 8 | 3.826 | 2.333 | 0.0227 | 1 |
| Snx18 | sorting nexin 18 | 25.506 | 0.809 | 0.0234 | 1 |
| Sntb1 | syntrophin beta 1 | 2.062 | 3.176 | 0.0241 | 1 |
| Ttll5 | tubulin tyrosine ligase like 5 | 16.106 | 0.98 | 0.0253 | 1 |
| Rpl21-ps14 | ribosomal protein L21, pseudogene 14 | 4.556 | 1.919 | 0.0257 | 1 |
| Prr22 | proline rich 22 | 2.286 | 2.954 | 0.0264 | 1 |
| Zfp335 | zinc finger protein 335 | 17.041 | 0.932 | 0.0264 | 1 |
| Srebf1 | sterol regulatory element binding transcription factor 1 | 32.17 | 0.719 | 0.0267 | 1 |
| Ccdc28b | coiled-coil domain containing 28B | 5.444 | 1.596 | 0.0281 | 1 |
| Fam118a | family with sequence similarity 118 member A | 29.637 | 0.677 | 0.0281 | 1 |
| Ptpn11 | protein tyrosine phosphatase non-receptor type 11 | 37.995 | 0.61 | 0.0283 | 1 |
| Cebpg | CCAAT enhancer binding protein gamma | 31.007 | 0.706 | 0.0286 | 1 |
| BC106179 | cDNA sequence BC106179 | 14.618 | 0.996 | 0.0292 | 1 |
| Zswim3 | zinc finger SWIM-type containing 3 | 6.079 | 1.485 | 0.0298 | 1 |
| Accs | 1-aminocyclopropane-1-carboxylate synthase homolog (inactive) | 20.297 | 0.851 | 0.0304 | 1 |
| Ctu1 | cytosolic thiouridylase subunit 1 | 14.644 | 0.944 | 0.0312 | 1 |
| Gm26566 | predicted gene, 26566 | 1.123 | 3.848 | 0.0322 | 1 |
| Ints8 | integrator complex subunit 8 | 14.105 | 1.003 | 0.0323 | 1 |
| Ube2s | ubiquitin conjugating enzyme E2 S | 2.266 | 2.647 | 0.0328 | 1 |
| Abcb7 | ATP binding cassette subfamily B member 7 | 15.259 | 0.948 | 0.0329 | 1 |
| Sorbs1 | sorbin and SH3 domain containing 1 | 1.672 | 3.479 | 0.033 | 1 |
| Rab5a | RAB5A, member RAS oncogene family | 24.447 | 0.747 | 0.0331 | 1 |
| Ddhd1 | DDHD domain containing 1 | 12.439 | 1.001 | 0.0334 | 1 |
| Socs6 | suppressor of cytokine signaling 6 | 18.949 | 0.817 | 0.0334 | 1 |
| Gm4705 | ribosomal protein L34 | 1.749 | 2.978 | 0.034 | 1 |
| Irf6 | interferon regulatory factor 6 | 17.451 | 0.87 | 0.0342 | 1 |
| Lrig1 | leucine rich repeats and immunoglobulin like domains 1 | 15.415 | 0.925 | 0.0343 | 1 |
| Echs1 | enoyl-CoA hydratase, short chain 1 | 23.411 | 0.723 | 0.0352 | 1 |
| Irf2bp1 | interferon regulatory factor 2 binding protein 1 | 41.727 | 0.594 | 0.0352 | 1 |
| Adam8 | ADAM metallopeptidase domain 8 | 1.477 | 3.282 | 0.0354 | 1 |
| Ddx1 | DEAD-box helicase 1 | 25.007 | 0.757 | 0.0356 | 1 |
| Hsph1 | heat shock protein family H (Hsp110) member 1 | 25.175 | 0.702 | 0.0356 | 1 |
| Tmem161b | transmembrane protein 161B | 23.533 | 0.772 | 0.0361 | 1 |
| Tanc1 | tetratricopeptide repeat, ankyrin repeat and coiled-coil containing 1 | 33.792 | 0.625 | 0.0361 | 1 |
| Gpr34 | G protein-coupled receptor 34 | 7.85 | 1.286 | 0.0362 | 1 |
| Heatr5b | HEAT repeat containing 5B | 12.002 | 1.068 | 0.0363 | 1 |
| Elovl7 | ELOVL fatty acid elongase 7 | 5.397 | 1.571 | 0.0365 | 1 |
| Mcat | malonyl-CoA-acyl carrier protein transacylase | 10.581 | 1.211 | 0.0375 | 1 |
| Cers6 | ceramide synthase 6 | 21.498 | 0.792 | 0.0376 | 1 |

| | | | | | |
|---|---|---|---|---|---|
| Simc1 | SUMO interacting motifs containing 1 | 20.655 | 0.788 | 0.038 | 1 |
| Nacc2 | NACC family member 2 | 28.274 | 0.675 | 0.0381 | 1 |
| S100a4 | S100 calcium binding protein A4 | 6.935 | 1.629 | 0.0392 | 1 |
| Ube2t | ubiquitin conjugating enzyme E2 T | 6.31 | 1.573 | 0.0392 | 1 |
| Gm17251 | predicted gene, 17251 | 8.589 | 1.257 | 0.0395 | 1 |
| Leng9 | leukocyte receptor cluster member 9 | 5.619 | 1.551 | 0.0402 | 1 |
| Arl8b | ADP ribosylation factor like GTPase 8B | 30.516 | 0.602 | 0.0407 | 1 |
| Itgae | integrin subunit alpha E | 1.058 | 3.742 | 0.041 | 1 |
| Tirap | TIR domain containing adaptor protein | 4.001 | 2.052 | 0.0415 | 1 |
| Pigk | phosphatidylinositol glycan anchor biosynthesis class K | 19.111 | 0.777 | 0.0419 | 1 |
| Otud1 | OTU deubiquitinase 1 | 32.017 | 0.659 | 0.0428 | 1 |
| Gm12115 | predicted gene 12115 | 4.019 | 1.718 | 0.0431 | 1 |
| Rgs1 | regulator of G protein signaling 1 | 21.004 | 0.821 | 0.0435 | 1 |
| Gzmm | granzyme M | 1.372 | 3.277 | 0.0436 | 1 |
| Mfsd11 | major facilitator superfamily domain containing 11 | 8.119 | 1.316 | 0.0443 | 1 |
| Rnf34 | ring finger protein 34 | 23.887 | 0.701 | 0.045 | 1 |
| Sec24d | SEC24 homolog D, COPII coat complex component | 3.238 | 2.085 | 0.0451 | 1 |
| Gpr18 | G protein-coupled receptor 18 | 38.907 | 0.587 | 0.0451 | 1 |
| A830010M20Rik | KIAA1107 | 9.998 | 1.145 | 0.0456 | 1 |
| Mcm3ap | minichromosome maintenance complex component 3 associated protein | 52.48 | 0.507 | 0.0456 | 1 |
| Dgkh | diacylglycerol kinase eta | 2.312 | 2.339 | 0.0459 | 1 |
| Hid1 | HID1 domain containing | 37.959 | 0.559 | 0.0463 | 1 |
| Akr7a5 | aldo-keto reductase family 7 member A2 | 16.801 | 0.784 | 0.0468 | 1 |
| Hddc2 | HD domain containing 2 | 24.907 | 0.697 | 0.0468 | 1 |
| Dpp9 | dipeptidyl peptidase Dpp9 | 34.488 | 0.635 | 0.0469 | 1 |
| Nme4 | NME/NM23 nucleoside diphosphate | 8.116 | 1.192 | 0.0472 | 1 |
| Gm12250 | predicted gene 12250 | 13.351 | 0.991 | 0.0473 | 1 |
| D930016D06Rik | Riken cDNA D930016D06 gene | 16.165 | 0.811 | 0.0477 | 1 |
| Tyrobp | TYRO protein tyrosine kinase binding | 1.006 | 3.684 | 0.0479 | 1 |
| F2rl2 | F2R like trypsin receptor 2 | 49.369 | 0.855 | 0.0479 | 1 |
| Ing5 | inhibtor of growth family member 5 | 62.657 | 0.526 | 0.048 | 1 |
| Rffl | ring finger and FYVE like domain | 11.468 | 0.963 | 0.0481 | 1 |
| App | amyloid beta precursor protein | 29.145 | 0.632 | 0.0481 | 1 |
| Ckap5 | cytoskeleton associated protein 5 | 38.678 | 0.595 | 0.0481 | 1 |
| Naip5 | NLR family, apoptosis inhibitory protein | 0.988 | 3.668 | 0.0485 | 1 |
| Ctsw | cathepsin W | 27.174 | 1.326 | 0.0487 | 1 |
| Gm6245 | UBX domain protein 2A pseudogene | 0.923 | 3.548 | 0.0489 | 1 |
| Casp3 | caspase 3 | 7.17 | 1.361 | 0.049 | 1 |
| **Downregulated genes** | | | | | |
| **Symbol** | **Entrez Gene Name** | **Basemean** | **Log2(Fold change)** | **Pvalue** | **Padj** |
| Nabp2 | nucleic acid binding protein 2 | 13.881 | -1.77 | 0.000322 | 0.766 |
| Rdh13 | retinol dehydrogenase 13 | 8.624 | -2.201 | 0.00131 | 1 |
| Arf5 | ADP ribosylation factor 5 | 132.093 | -0.575 | 0.00172 | 1 |
| 1700066M21Rik | chromosome 2 open reading frame 69 | 5.375 | -2.522 | 0.00338 | 1 |
| Ern1 | endoplasmic reticulum to nucleus signaling 1 | 2.712 | -3.926 | 0.0035 | 1 |
| Mdc1 | mediator of DNA damage checkpoint 1 | 43.003 | -0.798 | 0.00362 | 1 |
| Ythdf1 | YTH N6-methyladenosine RNA binding protein 1 | 72.9 | -0.624 | 0.00365 | 1 |
| Nfkbil1 | NFKB inhibitor like 1 | 3.954 | -3.206 | 0.00376 | 1 |
| 9930012K11Rik | chromosome 8 open reading frame 58 | 9.432 | -1.633 | 0.0047 | 1 |
| Cisd3 | CDGSH iron sulfur domain 3 | 37.352 | -0.803 | 0.00479 | 1 |
| Csrp1 | cysteine and glycine rich protein 1 | 54.059 | -0.707 | 0.00499 | 1 |

| | | | | | |
|---|---|---|---|---|---|
| Nedd4l | NEDD4 like E3 ubiquitin protein ligase | 26.802 | -1.036 | 0.00532 | 1 |
| Fam207a | family with sequence similarity 207 member A | 33.652 | -0.8 | 0.00654 | 1 |
| Pcbp1 | poly(rC) binding protein 1 | 99.487 | -0.512 | 0.00695 | 1 |
| Pkd1 | polycystin 1, transient receptor potential channel interacting | 40.395 | -0.71 | 0.0077 | 1 |
| Prickle1 | prickle planar cell polarity protein 1 | 4.05 | -2.633 | 0.00867 | 1 |
| Rbm10 | RNA binding motif protein 10 | 57.713 | -0.653 | 0.00889 | 1 |
| Alkbh7 | alkB homolog 7 | 14.495 | -1.186 | 0.00969 | 1 |
| Cbx6 | chromobox 6 | 49.47 | -0.692 | 0.00974 | 1 |
| Morc2a | MORC family CW-type zinc finger 2 | 14.151 | -1.231 | 0.0103 | 1 |
| Txlna | taxilin alpha | 60.781 | -0.615 | 0.0105 | 1 |
| Fig4 | FIG4 phosphoinositide 5-phosphatase | 9.428 | -1.405 | 0.0117 | 1 |
| Acsf2 | acyl-CoA synthetase family member 2 | 1.493 | -3.903 | 0.012 | 1 |
| Rab11fip3 | RAB11 family interacting protein 3 | 5.498 | -2.023 | 0.0121 | 1 |
| Chst14 | carbohydrate sulfotransferase 14 | 3.616 | -2.846 | 0.0122 | 1 |
| St3gal1 | ST3 beta-galactoside alpha-2,3-sialyltransferase 1 | 64.863 | -0.554 | 0.0133 | 1 |
| Slc16a5 | solute carrier family 16 member 5 | 27.127 | -0.809 | 0.0139 | 1 |
| Dcp2 | decapping mRNA 2 | 52.402 | -0.624 | 0.014 | 1 |
| Trim33 | tripartite motif containing 33 | 53.022 | -0.766 | 0.0142 | 1 |
| 9930111J21Rik1 | predicted gene 12185 | 1.443 | -3.817 | 0.0147 | 1 |
| Srf | serum response factor | 64.756 | -0.53 | 0.0149 | 1 |
| Gm5871 | predicted gene 5871 | 8.057 | -1.668 | 0.0153 | 1 |
| Thtpa | thiamine triphosphatase | 8.735 | -1.42 | 0.0155 | 1 |
| Lair1 | leukocyte associated immunoglobulin like receptor 1 | 8.076 | -1.667 | 0.0188 | 1 |
| Cst7 | cystatin F | 7 | -1.585 | 0.0194 | 1 |
| Exosc3 | exosome component 3 | 7.989 | -1.425 | 0.021 | 1 |
| Pogz | pogo transposable element derived with ZNF domain | 46.004 | -0.582 | 0.0212 | 1 |
| Dbt | dihydrolipoamide branched chain transacylase E2 | 22.22 | -0.823 | 0.023 | 1 |
| Atf6b | activating transcription factor 6 beta | 37.232 | -0.65 | 0.0235 | 1 |
| Mon1a | MON1 homolog A, secretory trafficking associated | 12.962 | -1.113 | 0.0235 | 1 |
| Coa7 | cytochrome c oxidase assembly factor 7 (putative) | 20.908 | -0.862 | 0.0237 | 1 |
| Mif | macrophage migration inhibitory factor | 32.073 | -0.697 | 0.0247 | 1 |
| Pfn2 | profilin 2 | 3.313 | -2.456 | 0.0247 | 1 |
| Gm10138 | predicted gene 10138 | 6.913 | -1.528 | 0.0252 | 1 |
| Smg5 | SMG5 nonsense mediated mRNA decay factor | 37.49 | -0.616 | 0.0262 | 1 |
| Xlr3b | X-linked lymphocyte-regulated 3C | 13.638 | -1.056 | 0.0276 | 1 |
| Bax | BCL2 associated X, apoptosis regulator | 3.791 | -2.243 | 0.0284 | 1 |
| D11Wsu47e | chromosome 17 open reading frame 80 | 5.172 | -1.675 | 0.0285 | 1 |
| Dars | aspartyl-tRNA synthetase 1 | 44.51 | -0.583 | 0.0287 | 1 |
| Vaultrc5 | vault RNA component 5 | 6.943 | -1.418 | 0.0289 | 1 |
| Gm20492 | predicted gene 20492 | 6.145 | -1.645 | 0.0299 | 1 |
| Ruvbl2 | RuvB like AAA ATPase 2 | 4.547 | -2.142 | 0.0299 | 1 |
| Rpp21 | ribonuclease P/MRP subunit p21 | 54.207 | -0.503 | 0.03 | 1 |
| Acap2 | ArfGAP with coiled-coil, ankyrin repeat and PH domains 2 | 44.434 | -0.597 | 0.0304 | 1 |
| B3gnt9 | UDP-GlcNAc:betaGal beta-1,3-N-acetylglucosaminyltransferase 9 | 1.323 | -3.726 | 0.0305 | 1 |
| Afg3l2 | AFG3 like matrix AAA peptidase subunit 2 | 22.388 | -0.862 | 0.0325 | 1 |
| 4933433G15Rik | RIKEN cDNA 4933433G15 gene | 1.222 | -3.552 | 0.0337 | 1 |
| Malsu1 | mitochondrial assembly of ribosomal large subunit 1 | 26.059 | -0.723 | 0.0344 | 1 |
| Pinx1 | PIN2 (TERF1) interacting telomerase inhibitor 1 | 28.205 | -0.678 | 0.0346 | 1 |
| Gm9385 | predicted pseudogene 9385 | 11.165 | -1.162 | 0.0346 | 1 |
| Rbpj | recombination signal binding protein for immunoglobulin kappa J region | 34.68 | -0.628 | 0.0349 | 1 |
| U2af1 | U2 small nuclear ribonucleoprotein auxiliary factor (U2AF) 1 | 24.779 | -0.78 | 0.0351 | 1 |

| | | | | | |
|---|---|---|---|---|---|
| Gm7266 | predicted gene 7266 | 5.952 | -1.53 | 0.0367 | 1 |
| Alas1 | 5'-aminolevulinate synthase 1 | 12.757 | -1.068 | 0.0368 | 1 |
| Dpf2 | double PHD fingers 2 | 35.936 | -0.599 | 0.0369 | 1 |
| Pbx4 | PBX homeobox 4 | 2.296 | -2.848 | 0.0371 | 1 |
| Tdp1 | tyrosyl-DNA phosphodiesterase 1 | 16.609 | -0.875 | 0.0374 | 1 |
| Ecd | ecdysoneless cell cycle regulator | 39.575 | -0.57 | 0.0378 | 1 |
| Fastkd5 | FAST kinase domains 5 | 4.298 | -1.866 | 0.038 | 1 |
| Nvl | nuclear VCP like | 49.236 | -0.505 | 0.0398 | 1 |
| Grm6 | glutamate metabotropic receptor 6 | 1.595 | -3.085 | 0.0403 | 1 |
| Chmp1a | charged multivesicular body protein 1A | 43.575 | -0.544 | 0.0405 | 1 |
| Dctpp1 | dCTP pyrophosphatase 1 | 12.008 | -1.115 | 0.0405 | 1 |
| Myo1c | myosin IC | 4.829 | -1.704 | 0.0405 | 1 |
| Mrpl21 | mitochondrial ribosomal protein L21 | 40.593 | -0.543 | 0.0407 | 1 |
| Capza1 | capping actin protein of muscle Z-line subunit alpha 1 | 17.883 | -0.816 | 0.0411 | 1 |
| Mrm1 | mitochondrial rRNA methyltransferase 1 | 24.754 | -0.776 | 0.0412 | 1 |
| Abr | ABR activator of RhoGEF and GTPase | 18.204 | -0.809 | 0.0414 | 1 |
| Abhd11 | abhydrolase domain containing 11 | 27.659 | -0.664 | 0.0418 | 1 |
| Isoc2a | isochorismatase domain containing 2 | 19.441 | -0.768 | 0.042 | 1 |
| Tmem156 | transmembrane protein 156 | 21.552 | -0.763 | 0.0422 | 1 |
| Chac2 | ChaC cation transport regulator homolog 2 | 1.5 | -3.085 | 0.0428 | 1 |
| Camk2a | calcium/calmodulin dependent protein kinase II alpha | 1.573 | -3.025 | 0.0433 | 1 |
| Eif2b5 | eukaryotic translation initiation factor 2B subunit epsilon | 38.404 | -0.593 | 0.0438 | 1 |
| Gypc | glycophorin C (Gerbich blood group) | 13.663 | -0.976 | 0.0438 | 1 |
| Swi5 | SWI5 homologous recombination repair protein | 42.052 | -0.564 | 0.0442 | 1 |
| Slc7a6os | solute carrier family 7 member 6 opposite strand | 25.042 | -0.685 | 0.0442 | 1 |
| Wdr83os | WD repeat domain 83 opposite strand | 19.831 | -0.826 | 0.0444 | 1 |
| Krit1 | KRIT1 ankyrin repeat containing | 32.123 | -0.604 | 0.0446 | 1 |
| Gpr89 | G protein-coupled receptor 89B | 20.725 | -0.816 | 0.0448 | 1 |
| Unc45a | unc-45 myosin chaperone A | 27.474 | -0.671 | 0.0449 | 1 |
| Cdr2 | cerebellar degeneration related protein 2 | 10.757 | -1.129 | 0.045 | 1 |
| Fancb | FA complementation group B | 1.142 | -3.408 | 0.0455 | 1 |
| Gm13623 | 60S ribosomal protein L13-like | 1.774 | -3.194 | 0.0464 | 1 |
| Dpp8 | dipeptidyl peptidase 8 | 32.959 | -0.585 | 0.0472 | 1 |
| Klhl17 | kelch like family member 17 | 9.92 | -1.083 | 0.0476 | 1 |
| E230001N04Rik | RIKEN cDNA E230001N04 gene | 9.284 | -1.193 | 0.048 | 1 |
| Gm4034 | predicted gene 4034 | 1.629 | -3.11 | 0.0483 | 1 |
| Itfg2 | integrin alpha FG-GAP repeat containing 2 | 19.651 | -0.797 | 0.0485 | 1 |
| Pop1 | POP1 homolog, ribonuclease P/MRP subunit | 13.937 | -0.891 | 0.0496 | 1 |

### Supplementary Figures

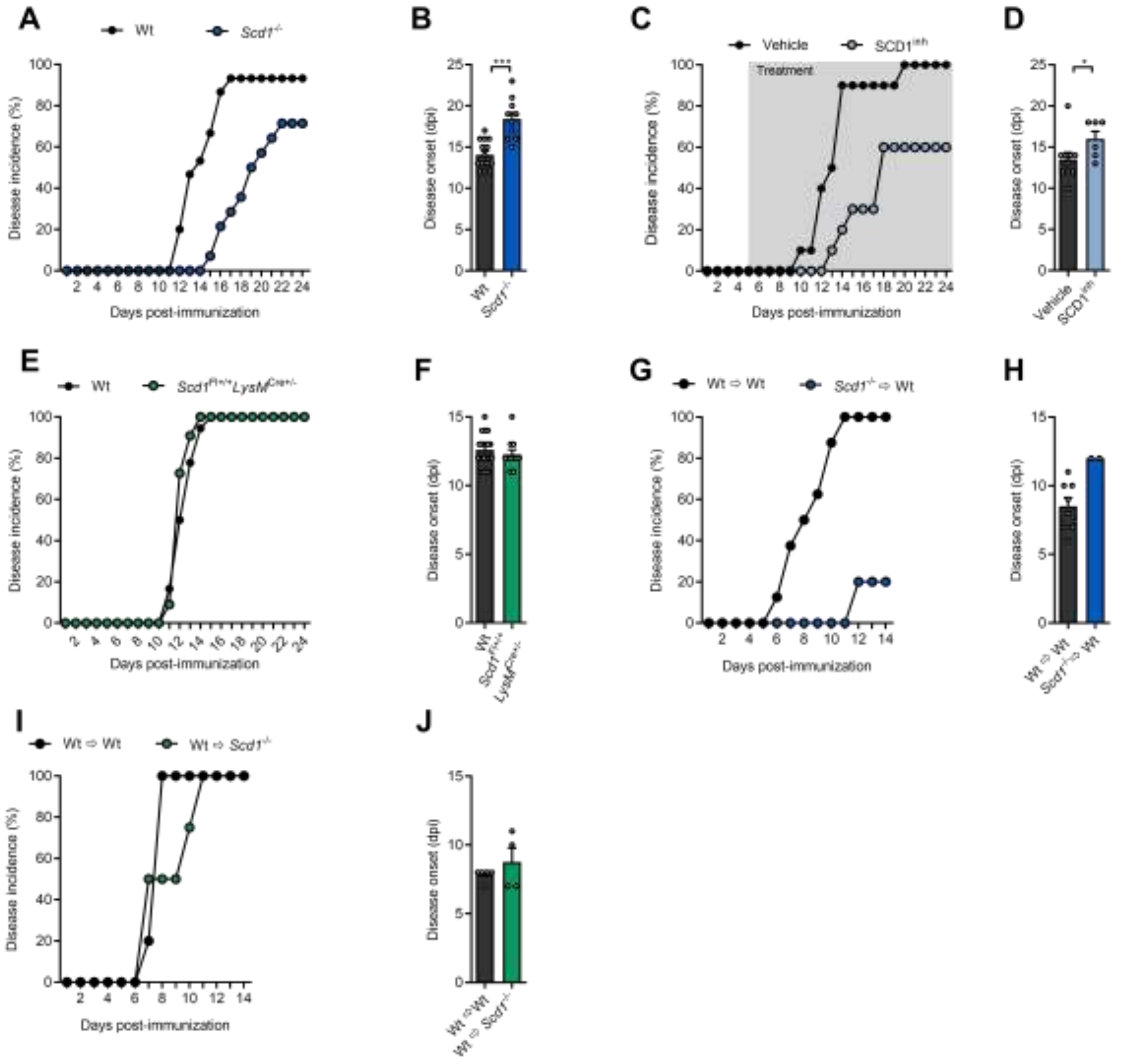


**Supplemental Fig. 1. Pharmacological inhibition and genetic deficiency of SCD1 reduces EAE disease severity.** EAE disease incidence (A,C,E,G,I) and onset (B,D,F,H,J) of wild-type (wt, n=15) and *Scd1*$^{-/-}$ (n=14) mice (A,B), vehicle- (n=10) and SCD1 inhibitor-treated (SCD1$^{inh}$, 2.5 mg/kg, n=10) mice (C,D), wt (n=19) and *Scd1*$^{Fl+/+}$ *LysM*$^{Cre+/-}$ (n=11) mice (E,F), wt mice that received wt (Wt ⇨ Wt, n=8) or *Scd1*$^{-/-}$ (*Scd1*$^{-/-}$ ⇨ Wt, n=10) encephalitogenic lymphocytes (G,H), wt (n=5) and *Scd1*$^{-/-}$ (n=4) mice that received wt encephalitogenic lymphocytes (I,J). Animals that did not develop EAE were not included in the disease onset analysis. Dpi: days post-immunization. All replicates were biologically independent. All data are represented as mean ± SEM. *, $P < 0.05$; **, $P<0.01$; ***, $P < 0.001$; calculated with two-tailed unpaired student T-test (A-G,I,J), or Mann-Whitney analysis (H).

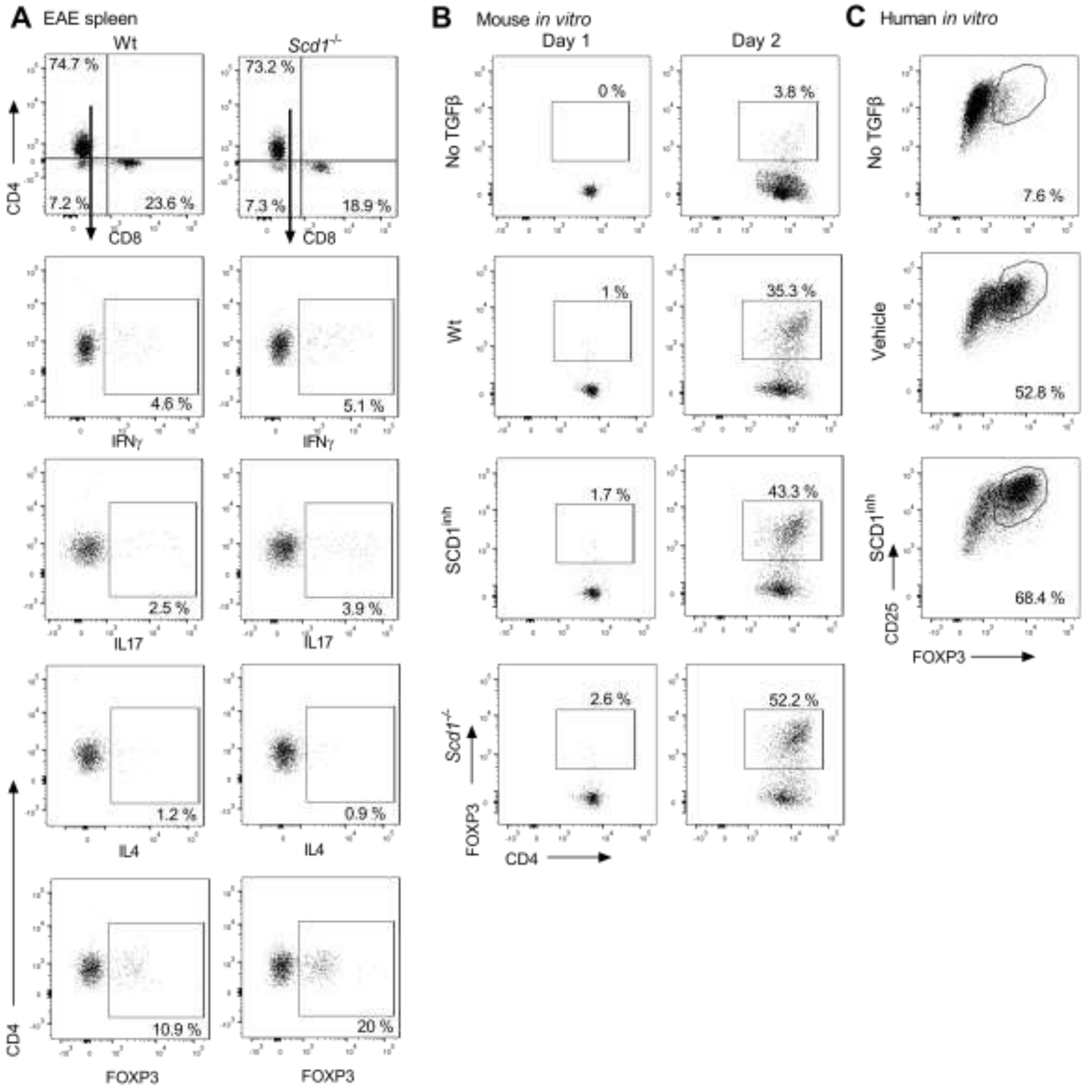


**Supplemental Fig. 2. *Scd1* deficiency increases Treg differentiation. (A)** Frequency of CD4$^+$, CD8$^+$, CD4$^-$CD8$^-$, CD4$^+$IFNγ$^+$, CD4$^+$IL17$^+$, CD4$^+$IL4$^+$, and CD4$^+$FOXP3$^+$ in the spleen of wild-type (wt, n=11 animals) and *Scd1*$^{-/-}$ (n=11 animals) EAE animals 10 days after EAE induction. Representative flow cytometric plots are shown. **(B,C)** Representative flow cytometric plots of wt and *Scd1*$^{-/-}$ mouse naïve T cells and human naïve T cells differentiated under Treg polarizing conditions and treated with vehicle or SCD1 inhibitor (SCD1$^{inh}$, CAY10566, 1 μM). For mouse T cell cultures, the frequency of CD4$^+$FOXP3$^+$ cells was quantified one and two days after Treg induction. For human T cell cultures, frequency of CD25$^{Hi}$FOXP3$^+$ cells was quantified four days after induction.

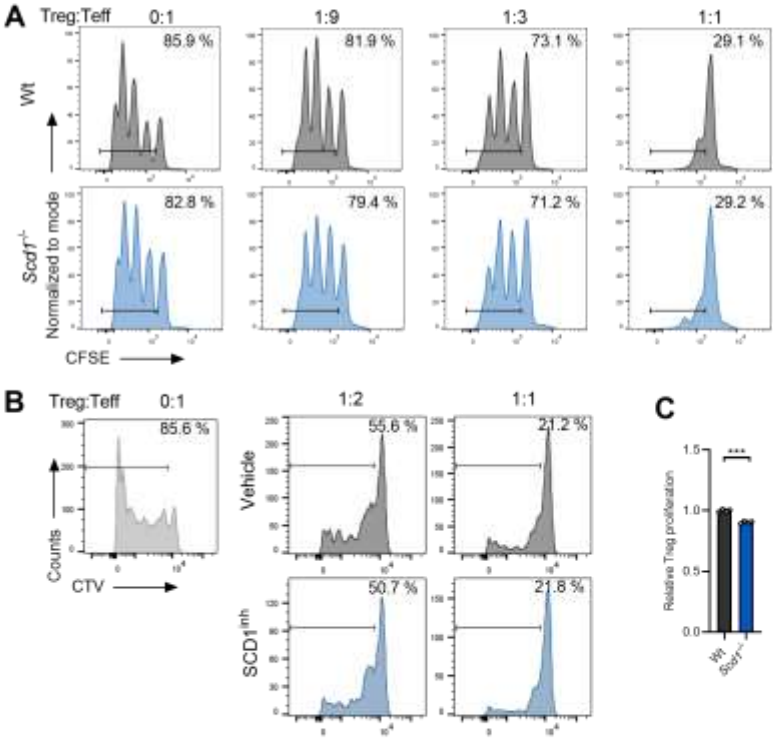


**Supplemental Fig. 3. Loss of *Scd1* does not affect the suppressive and proliferative capacity of regulatory T cells. (A,B)** Representative flow cytometric plots of the suppressive capacity of wild-type (wt) and *Scd1*$^{-/-}$ mouse Tregs (A), or human Tregs differentiated under the presence of SCD1 inhibitor (SCD1$^{inh}$, CAY10566, 1 μM) or vehicle (B). Increasing amounts of Tregs were cultured with CFSE- or CellTrace Violet (CTV)-labeled CD4$^{+}$CD25$^{-}$ effector T cells (D, n=3 samples; E, n=2 healthy controls). Percentage proliferation was assessed after three (A) or five (B) days. **(C)** Mouse CD4$^{+}$CD25$^{+}$ regulatory T cells were labelled with CFSE, stimulated with anti-CD3ε (2 μg/ml), anti-CD28 (2 μg/ml), recombinant IL-2 (5 U/ml), and analyzed after four days with flow cytometry (n=4 samples). All data are represented as mean ± SEM. ***, $P < 0.001$; calculated with two-tailed unpaired student T-test.

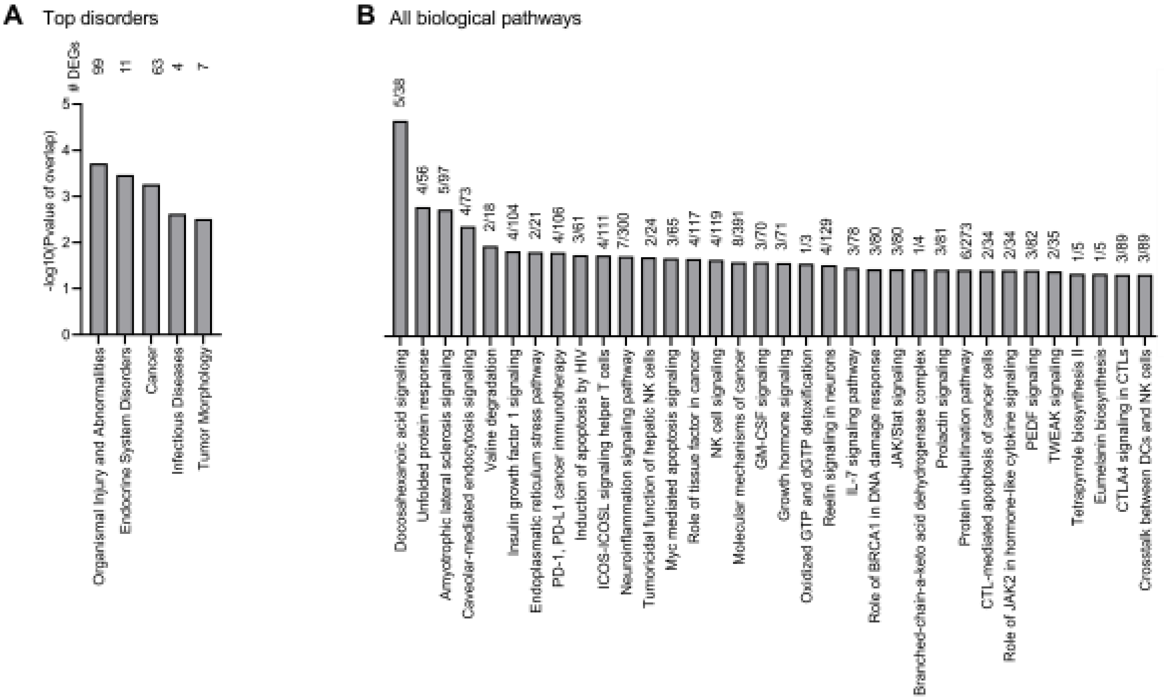


**Supplemental Fig. 4. Pathway analysis of differentially expressed genes in wt and *Scd1*$^{-/-}$ naïve T cells.** Bulk RNA sequencing was performed on wild-type (wt) and *Scd1*$^{-/-}$ naïve T cells. Disorders and canonical pathways associated with *Scd1*$^{-/-}$ T cells identified using Ingenuity Pathway Analysis. DEGs: differentially expressed genes. All results are pooled from four independent experiments.

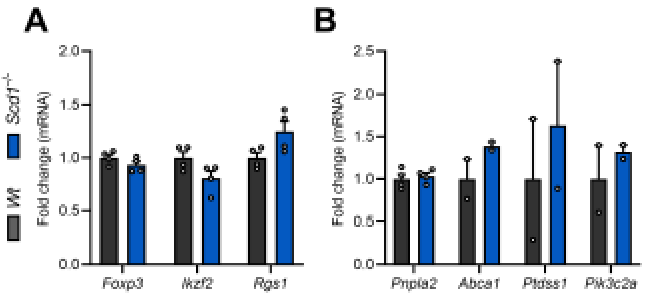


**Supplemental Fig. 5. *Scd1*-deficiency does not affect the transcriptional profile of regulatory T cells.** Expression of regulatory T cell-associated functional genes *Foxp3*, *Ikzf2*, and *Rgs1*, and DHA-signaling genes *Pnpla2*, *Abca1*, *Ptdss1*, and *Pik3c2a* in $CD4^+CD25^+$ regulatory T cells isolated from naive *Scd1*$^{-/-}$ mice and wild-type (wt) littermates (n=4 mice/group). Data are represented as mean ± SEM; calculated with two-tailed unpaired student T-test or Mann-Whitney analysis.

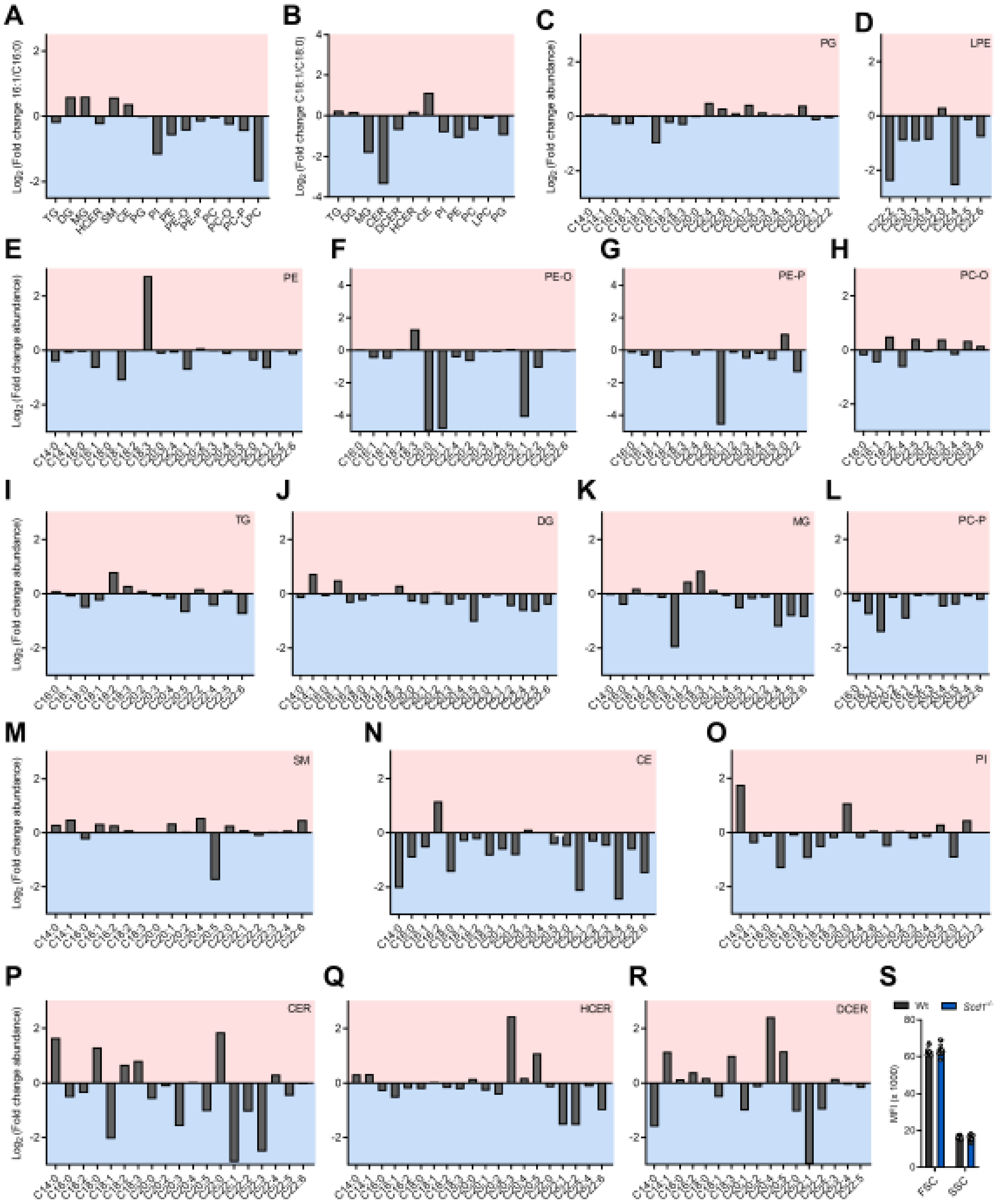


**Supplemental Fig. 6. *Scd1$^{-/-}$* naïve T cells show a decreased intracellular abundance of fatty acid-containing lipids. (A-R)** Liquid chromatography electrospray ionization tandem mass spectrometry (LC-ESI-MS/MS) analysis was used to define the lipidome of wild-type (wt) and *Scd1$^{-/-}$* naïve T cells (n=2 samples). $Log_2$ fold change abundance of fatty acid species within each lipid class is shown (*Scd1$^{-/-}$* vs. wt). Only detectable lipid species and fatty acyl moieties are shown. **(S)** Flow cytometric analysis of forward scatter (FSC) and sideward scatter (SSC) of wt and *Scd1$^{-/-}$* naïve

T cells. Results are pooled from two (A-R) or 12 (S) independent experiments. Data are represented as mean (A-R) or as mean ± SEM (S); calculated with two-tailed unpaired student T-test.

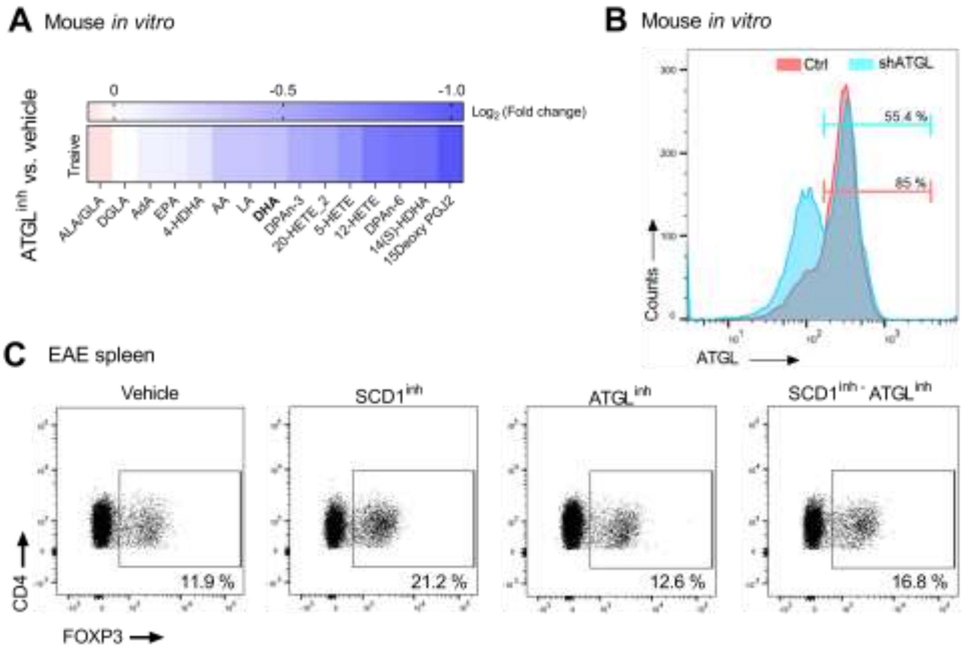


**Supplemental Fig. 7. ATGL inhibition decreases intracellular non-esterified DHA levels in naïve *Scd1*-deficient T cells. (A)** Liquid chromatography tandem mass spectrometry (LC-MS/MS) analysis was performed to define the abundance of intracellular non-esterified fatty acids and downstream metabolites in *Scd1*$^{-/-}$ naïve T cells treated with vehicle (n=2 samples) or ATGL inhibitor (ATGL$^{inh}$, Atglistatin, 20 µM; n=2 samples) for two days. Log$_2$ fold change abundance of all detectable fatty acids and downstream metabolites is shown (Atglistatin-treated vs. vehicle-treated). **(B)** ShRNA-mediated gene silencing of *Atgl* in mouse naïve CD4$^+$ T cells. **(C)** Representative flow cytometric plots of CD4$^+$FOXP3$^+$ cell frequency in the spleen of EAE mice treated with an SCD1 inhibitor (SCD1$^{inh}$, CAY10566, 2.5 mg/kg), ATGL inhibitor (ATGL$^{inh}$, Atglistatin, 200 µmol/kg), a combination, or vehicle (oral gavage twice a day with a 12 h interval). Cells were collected 10 days post-immunization. All results are pooled from two independent experiments. Data are represented as mean.

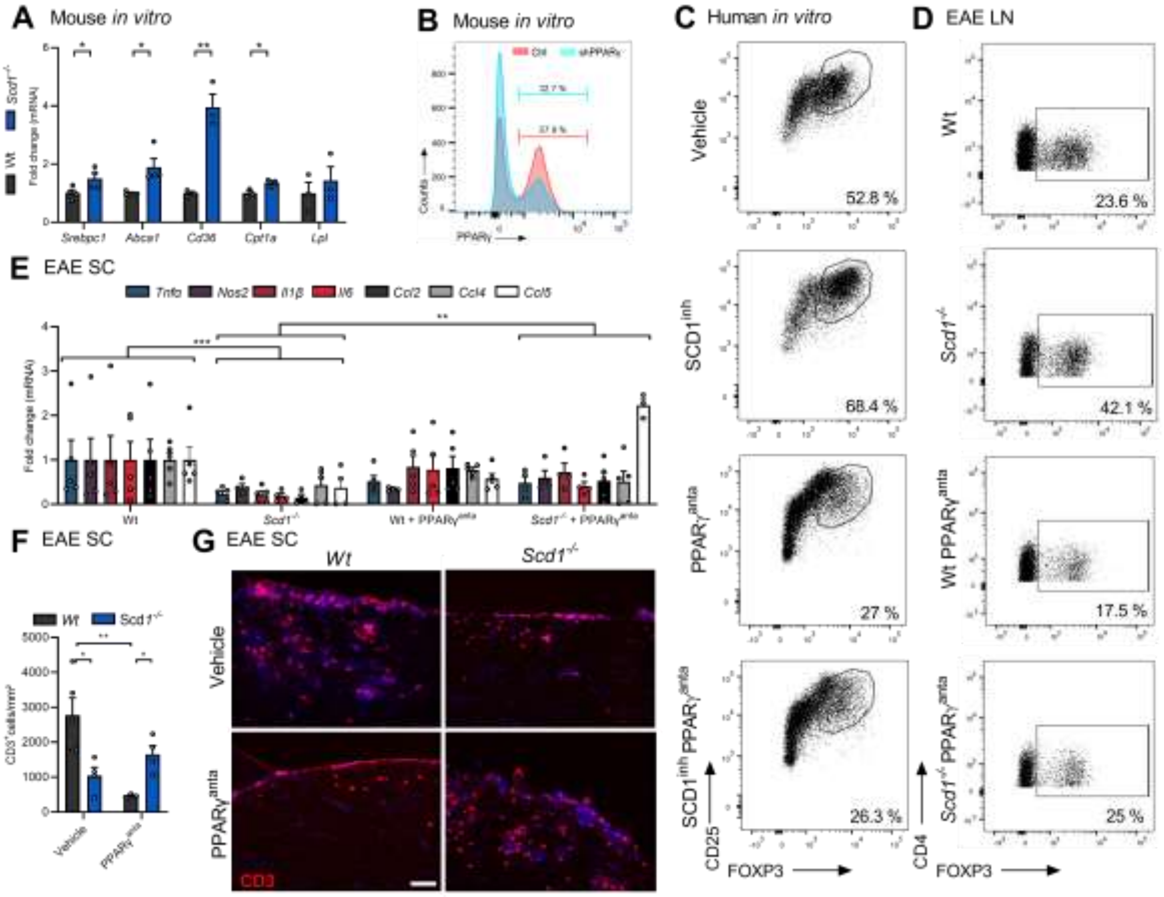


**Supplemental Fig. 8. PPARγ signaling promotes Treg differentiation in the absence of SCD1. (A)** mRNA expression of PPARγ-response genes *Srebpc1*, *Abca1*, *Cd36*, *Cpt1a* and *Lpl* in wt and *Scd1*$^{-/-}$ naïve T cells (n=3-5 samples). **(B)** ShRNA-mediated gene silencing of *Pparγ* in naïve mouse CD4$^{+}$ T cells. **(C)** Human naïve T cells were differentiated under Treg polarizing conditions and treated with vehicle (n=5 healthy controls) or PPARγ$^{anta}$ (25 μM; n=4 healthy controls). Frequency of CD4$^{+}$FOXP3$^{+}$ cells was quantified four days after Treg induction. Representative flow cytometric plots are shown. **(D-G)** Wt and *Scd1*$^{-/-}$ EAE animals were treated daily with a PPARγ-antagonist (GW9662, 2mg/kg, n=5/group) or vehicle, and were sacrificed 28 days after EAE induction. Frequency of CD4$^{+}$, CD8$^{+}$, CD4$^{-}$CD8$^{-}$, CD4$^{+}$IFNγ$^{+}$, CD4$^{+}$IL17$^{+}$, CD4$^{+}$IL4$^{+}$, and CD4$^{+}$FOXP3$^{+}$ cells in the lymph nodes (D). mRNA expression of *Nos2*, *Tnfa*, *Il1β*, *Il6*, *Ccl2*, *Ccl4, and Ccl5* in spinal cord (SC) tissue (E). Quantification (F) and representative images (G) of CD3 staining of SC tissue (F,G). Data are represented as mean ± SEM. *, P<0.05; **, P < 0.01; ***, P<0.001; calculated with unpaired student T-test (A) and Mann-Whitney U test (E,F).
